# ARGminer: A web platform for crowdsourcing-based curation of antibiotic resistance genes

**DOI:** 10.1101/274282

**Authors:** G. A. Arango-Argoty, G. K. P. Guron, E. Garner, M.V. Riquelme, L. S. Heath, A. Pruden, P. J. Vikesland, L. Zhang

## Abstract

Curation of antibiotic resistance gene (ARG) databases is labor intensive and requires expert knowledge to manually collect, correct, and/or annotate individual genes. Consequently, most existing ARG databases contain only a small number of ARGs (~5k genes) and updates to these databases tend to be infrequent, commonly requiring years for completion and often containing inconsistencies. Thus a new approach is needed to achieve a truly comprehensive ARG database while also maintaining a high level of accuracy. Here we propose a new web-based curation system, ARGminer, that supports the annotation and inspection of several key attributes of potential ARGs, including gene name, antibiotic category, resistance mechanism, evidence for mobility and occurrence in clinically-important bacterial strains. Here we employ crowdsourcing as a novel strategy to overcome limitations of manual curation and expand curation capacity towards achieving a truly comprehensive and perpetually up-to-date database. Further, machine learning is employed as a powerful means to validate database curation, drawing from natural language processing to infer correct and consistent nomenclature for each potential ARG. We develop and validate the crowdsourcing approach by comparing performances of multiple cohorts of curators with varying levels of expertise, demonstrating that ARGminer is a time and cost efficient means of achieving accurate ARG curation. We further demonstrate the reliability of a trust validation filter for rejecting input generated by spammers. Crowdsourcing was found to be as accurate as expert annotation, with an accuracy >90% for the annotation of a diverse test set of ARGs. The ARGminer public search platform and database is available at http://bench.cs.vt.edu/argminer.

## INTRODUCTION

Antimicrobial resistance (AMR) has been identified by the World Health Organization (WHO) as a major global health threat. It is projected that AMR will increase exponentially by 2050, leading to substantial human morbidity and mortality {O’Neill, 2016 #3009;Pires, 2017 #3100}. Therefore, swift action is required to enable enhanced monitoring and help tackle the spread of AMR, including: understanding the mechanisms controlling dissemination of antibiotic resistance genes (ARGs) via environmental sources and pathways {Pruden, 2013 #3101;Martínez, 2008 #3034;Bengtsson-Palme, 2017 #3103}, discovering novel ARGs before they are found to be problematic in the clinic {Berglund, 2017 #3094}, developing new computational strategies for ARG annotation {Arango-Argoty, 2017 #3077;Gibson, 2015 #3058;Lakin, 2017 #3083;Yang, 2016 #294;McArthur, 2013 #3016}, and expansion of current ARG repositories {Lakin, 2017 #3083;Arango-Argoty, 2017 #3077}.

Metagenomic sequencing has provided a powerful means for accessing the diverse array of ARGs, or “resistomes,” {Sello, 2012 #3125} characteristic of various environments {Li, 2017 #3106;Garner, 2016 #3107;Bengtsson-Palme, 2016 #3108;Pal, 2016 #755} and has supported the discovery of novel ARGs and their interactions {Pehrsson, 2016 #3092;Forsberg, 2014 #3062}. Existing metagenomic approaches are largely dependent upon predicting antibiotic resistance attributes through sequence similarity computation, which is subject to major limitations. First, such similarity computations require a high quality and up-to-date ARG reference/annotation database to enable consistent and accurate ARG identification. Second, the scope of such analyses is limited to previously characterized ARGs, either due to the parameter cutoff stringency employed in the sequence alignment or to lack of a comprehensive target gene for alignment {Yang, 2016 #294}.

To improve the capacity of metagenomic-based approaches to broadly and accurately detect the full range of ARGs present in a given sample, it is necessary to continuously expand and improve curation of corresponding databases {Arango-Argoty, 2017 #3077}. However, risk of incorporation of false positives, i.e., “ARG-like” genes that do not necessarily induce an AMR phenotype, stands as a major impediment to expanded curation efforts. Therefore, manual inspection and validation of potential ARG entries is a critical aspect of ensuring the validity of AMR databases and their application.

Manual curation of ARGs is typically carried out by a few experts associated with research groups committed to maintaining public databases. This process is complex, tedious, and time-consuming. For instance, the last update of the Antibiotic Resistance Database (ARDB) was in 2009 {Liu, 2008 #3081}, and, therefore, it does not contain any newly discovered ARGs, such as *bla*_NDM-1_ or mcr-1. The MEGARes database {Lakin, 2017 #3083}, which was designed to simplify the organization of ARG annotation, has not been updated since December, 2016. The resqu database, which contains genes for which there is evidence of having been transferred via Mobile Genetic elements (MGEs), has not been updated since 2013 {Bengtsson-Palme, 2017 #3095}. The Comprehensive Antibiotic Resistance Database (CARD) {McArthur, 2013 #3016} is widely considered to be the most up-to-date ARG resource. First introduced in 2016, CARD has been updated a total of 21 times since its most recent release in October 2018, with corresponding changes to the ARG sequences and metadata (e.g., antibiotic class, gene name, and mechanism). This acutely illustrates how complex and time-consuming ARG database curation is, even for domain experts.

Attempts have been made to address limitations of currently-available databases, including introduction of new databases, such as the structured antibiotic resistance database (SARG), which employed intense manual curation to address issues such as inconsistencies in nomenclature and elimination of single-nucleotide polymorphisms and housekeeping genes {Yang, 2016 #3267}. In our own research group, we previously introduced DeepARG: a computational approach for predicting of ARGs using deep learning {Arango-Argoty, 2017 #3077}. Along with the machine learning models, we also released a curated database named DeepARG-DB. This database employs manual curation, literature review of ARGs, and annotation of ARGs using sequence alignments. DeepARG-DB was first released in July, 2017 and most recently updated August, 2018. However, deepARG database depends of annotations from multiple resources, making it sensitive to the propagation of errors from other databases. This highlights the need of enabling a specialized tool that brings all the ARG information from different resources to easily integrate new ARGs or to validate the annotations of current ARGs.

To overcome the difficulties in curation and manual validation of an extensive number of ARGs, a novel approach that breaks down this complex task into simpler and smaller microtasks is proposed. The core of this methodology consists of aggregating a compendium of AMR resources and deploying a crowdsourcing strategy, which simplifies the ARG information to allow nonexperts, i.e., the general public, and domain experts collectively to execute curation of the ARG database. Application of crowdsourcing in biology, particularly for data curation, is not new and comprises a variety of areas including: name entity recognition (NER) for drug and diseases {Islamaj Dogan, 2009 #3110;Khare, 2015 #3111;Lu, 2009 #3112}, identification of medically-relevant terms from patient online posts {MacLean, 2013 #3113}, annotation of diseases described in PubMed {Good, 2014 #3114}, and systematic examination of databases and other resources for drug indications, biomedical ontologies, and gene-disease interactions {Wei, 2013 #3115;Wei, 2012 #3116;Arighi, 2013 #3117;Lu, 2012 #3118;Khare, 2015 #3111}. Interestingly, in most of the studies, crowdsourcing has proven to be as effective as expert curation {Khare, 2015 #3111;Good, 2013 #3122}.

A major problem that encompasses all ARG resources is the lack of a standardized gene nomenclature. In particular, naming ARGs does not follow the general nomenclature for naming bacterial genes {Demerec, 1966 #3261}. For instance, macrolide resistance genes are structured so that the class is indicated by brackets (e.g., *ole*(B), *srm*(B), *vga(B)* or *ere*(B)) {Roberts, 1999 #3262}. When compared to tetracycline genes, this gene nomenclature differs radically, because, in tetracycline genes, the determinant is placed as a capital letter after the gene name (e.g., tetA, tetB, tetC) {Levy, 1999 #3263}. At the same time, those nomenclatures differ from the gene convention proposed to annotate beta lactamases genes {Hall, 2015 #3261}. Other examples to highlight these differences include the aminoglycoside gene {Vanhoof, 1998 #3264} aadA1 found under different names across the available ARG databases (ANT(3’’)-I, aadA1-pm, ANT3-DPRIME, and ant3ia). Therefore, the diversity and variation in the ARG nomenclature and naming conventions complicate and greatly hinder consistent ARG curation.

Here we introduce ARGminer, an online platform to enhance manual curation of ARGs. ARGminer enables users to curate and retrieve all the information available from several ARG resources, including CARD {McArthur, 2013 #3016}, DeepARG-DB {Arango-Argoty, 2017 #3077}, ARDB {Liu, 2008 #3081}, MEGARes {Lakin, 2017 #3083}, UniProt {Leplae, 2004 #3088}, the National Database of Antibiotic Resistant Organisms (NDARO) (https://www.ncbi.nlm.nih.gov/pathogens/antimicrobial-resistance/), the structured antibiotic resistance database (SARG) {Yang, 2016 #3267}, ResFinder {Zankari, 2012 #311}, and the ARGANNOT {Gupta, 2014 #308} databases. Manual crowd-source-based curation is enhanced by a machine learning model based on word embeddings {Goldberg, 2014 #3265;Turian, 2010 #3266}, a technique widely used in natural language processing (NLP) to aid in validation and achieve consistency in ARG nomenclature. On the other hand, mobile genetic elements (MGEs) such as plasmids phages and viruses play an important role in the dissemination of ARGs {Gillings, 2014 #3250; Leplae, 2004 #3088; Bengtsson-Palme, 2017 #3275}. Therefore, ARGminer also interfaces with the PATRIC {Wattam, 2014 #754} and Classification of Mobile Genetic Elements (ACLAME) {Leplae, 2004 #3088} databases, which provide information on potential carriage of ARGs by pathogens or MGEs, respectively. The ARGminer platform is designed, built, and implemented as an open-source project facilitating a collaborative and integrative approach for the standardization of ARG annotation by the broad community of scientists and citizens motivated by a common desire to contribute towards combating the spread of AMR. ARGminer also includes a community blog for users to post questions and share solutions/discussion regarding antimicrobial resistance with the objective to keep the scientific community actively engaged in the latest updates and development of ARG databases (see **Supplementary Figure 1**). All data associated with ARGminer, as well as the source code, is freely available under a public repository at http://bench.cs.vt.edu/argminer.

## MATERIALS AND METHODS

### ARG Database

ARGs were downloaded from the following resources: CARD {McArthur, 2013 #3016}, which contains ARG information; the ARDB {Liu, 2008 #3081} database, which comprises a vast number of homology-predicted ARGs; DeepARG-DB {Arango-Argoty, 2017 #3077}, which integrates ARGs from UniProt {Consortium, 2014 #3084}, CARD, and ARDB; MEGARes {Lakin, 2017 #3083} database, which incorporates genes from the ARG-ANNOT {Gupta, 2014 #308}, ResFinder {Zankari, 2012 #311}, the Lahey Clinic beta-lactamase archive {Bush, 2010 #3086} available from the National Center for Biotechnology Information (NCBI), the SARG database {Yang, 2016 #3267}, and the NDARO database version 2.

To obtain a clean collection of ARGs, the DeepARG-DB database was updated with a more recent version of the CARD (v 2.0.4) and UniProt databases using their corresponding sequence identifiers. Discontinued UniProt sequences were removed from DeepARG-DB, whereas the newly-added ARGs from CARD were incorporated. Also, genes from CARD known to confer resistance due to single point mutations were removed. All sequences from all databases were clustered to remove duplicates by using cd-hit and identity cutoff of 100%. The resulting collection of ARGs was then aligned to all databases using DIAMOND {Buchfink, 2015 #3174} and TBLASTN {Gertz, 2006 #3124} to extract the best hit of each ARG along with its corresponding metadata. In this manner, each ARG is represented by its best hit to each database, upholding consistency in annotation among the ARG resources. The metadata from the UniProt database is accessed via the UniProt API (Application Programming Interface), which allows retrieval of up-to-date information for each gene. Therefore, each ARG is displayed in the user interface as a set of sections containing an ARG’s best hits, its metadata, and the alignment quality. Scores are shown as bars to enhance readability and curator interpretation (see **Supplementary Figure 3A**).

### Mobile Genetic Element Curation

The ACLAME database {Leplae, 2004 #3088} primarily houses genes associated with plasmids, viruses, and phages and was used to identify ARGs that have potential of being mobilized by MGEs. DIAMOND {Buchfink, 2015 #3174} was used to compare ARGs to MGEs via sequence alignment (parameters e-value < 1e-10). Alignment information, along with MGE metadata, is presented in the interface for users to make a decision on whether an ARG has enough evidence of being carried by an MGE or not. This evidence is scored from 0 to 5. A colorimetric ranking scale depicts the degree of confidence for the information presented in the MGE panel (see **Supplementary Figure 3B**).

### Pathogen Sequence Curation

A total of 98,758 bacterial genomes were downloaded from the PATRIC {Wattam, 2014 #754} database. This database contains information about bacterial pathogenicity, antimicrobial resistance phenotype, corresponding diseases, and host organisms, information that is particularly valuable to clinicians seeking to identify pathogens and potential antibiotic resistance traits. For instance, the UniProt gene entry BAE06009.1 was present in 2,037 bacterial genomes, of which, 1,004 belong to pathogenic bacteria, 40 are involved in cystic fibrosis disease in humans, and 706 exhibit intermediate and resistant phenotypes (see **Supplementary Figure 3C**). The collection of ARGs were then screened against the genome sequences from PATRIC using DIAMOND {Buchfink, 2015 #3174}. To ensure the quality of the assignments, all genes with an identity below 90% and an alignment coverage below 90% were discarded. Users are asked to rate the potential pathogenicity of known bacterial hosts of ARGs based on the evidence provided by PATRIC (frequency of pathogenic genomes, diseases, antimicrobial phenotype, and hosts).

### Annotation microtasks

An annotation task consists of labeling ARGs based on the evidence provided on the web site. Users are requested to classify an ARG in terms of gene name, antibiotic class, and antibiotic mechanism. In addition, users are asked to rank the evidence of that this ARG sometimes occurs on MGEs or pathogens. A user-friendly web interface makes it easy to follow the annotation process (**Supplementary Figure 4**). Through crowdsourcing and converting the complex annotation tasks into achievable microtasks, ARGminer advocates mass collaboration from an open community that includes both experts and the general public to tackle the difficult task of ARG annotation.

### ARG nomenclature prediction

ARGminer includes a machine learning model based on a word embedding representation for the prediction of the gene name nomenclature given the text metadata information available from different databases. To this end, information such as ARG names, antibiotic classes, and other text data was collected from the CARD database {McArthur, 2013 #3016}. In total 2,355 ARGs’ metadata and names were used for training/testing the model. Labels were defined as the shape of the ARG names. For instance, the label for *opmE* is xxxX, the label for the tetracycline gene tet(A) is xxx(X), and the label for the Beta-lactamase gene TEM-21 is XXX-N. X, x and N correspond to a letter (uppercase or lowercase) and a number, respectively. The nomenclature data set for training and testing was built as follows:

1. Obtain ARG sequences, names, and metadata from CARD.
2. Align CARD sequences to other databases (deepARG-DB, ARDB, ResFinder, ARGANNOT, SARG, NDARO) and extract the best hit.
3. Extract corresponding metadata for the best hits.
4. Build the data set.

Once the data set was built, 80% of the entries were randomly selected for training and the remaining for validation. This process was repeated ten times to check consistency and variability of the results. FastText, a library for text classification and representation using word embeddings {Bojanowski, 2016 #3203; Joulin, 2016 #3078}, was used to build the model. Briefly, the model was trained with default parameters along an embedding space of 100 dimensions during 100 epochs. **Supplementary Figure 5** shows the workflow of the ARG nomenclature prediction framework.

### Expert gold standard data set

To assess the accuracy and quality of classifications generated by crowd-sourcing, three domain experts who are actively engaged in environmental ARG research were asked to annotate a gold standard set of 35 ARGs by name, antibiotic class, and mechanism. In total, 34 out of the 35 ARG annotations were in agreement among at least two of the three experts in terms of antibiotic class and gene name. These 34 ARGs were further considered in downstream analysis.

### Crowdsourcing microtasks

Annotations were performed by two groups of curators. One group was recruited via Amazon Mechanical Turk (MTurk), an online platform that allows access to a broad crowdsourcing audience to perform Human Intelligent Tasks (HITs). When a curator performs an annotation, ARGminer prompts a token that curators need to submit to the MTurk web site for validation to then obtain a monetary reward. Because of the high diversity of MTurk curator backgrounds, the ARGminer HITs were opened to a broad audience, including domain experts and nonexperts, and curators were allowed to perform a limited number of annotations (maximum 20). Another group of curators consisted of students enrolled in a graduate-level microbiology class. Not all of the students possessed deep antimicrobial resistance domain knowledge, but they all had general familiarity with microbiology.

### User interface

The ARGminer interface has three main components or sections:

1. **Current Annotation**: Summarizes current available information for a given ARG. It consists of the gene name, antibiotic class, database from which the sequence was extracted, and number of times the gene has been inspected by curators (see **Figure 1A**).
2. **Evidence**: Corresponds to the metadata available for the ARG as well as the best hit from the CARD, ARDB, and MEGARes databases. It also provides evidence/information regarding whether the gene is likely carried by an MGE (the ACLAME database) and whether the gene is known to be found in pathogen genomes (the PATRIC database, see **Figure 1B**).
3. **Microtask**: Refers to the section where a curator enters his/her annotation. The information in this panel must be consistent with the observations from the evidence. It consists of three simple steps. First, curators must validate the gene name, antibiotic class, and mechanism by looking at the Evidence section. Second, curators must rank the MGE and pathogen evidence. Third, curators must rank their overall annotation by scoring their expertise (how familiar they are with ARGs) and confidence (how strong the evidence is, see **Figure 1C and Supplementary Figure 4**).

**Figure 1:**
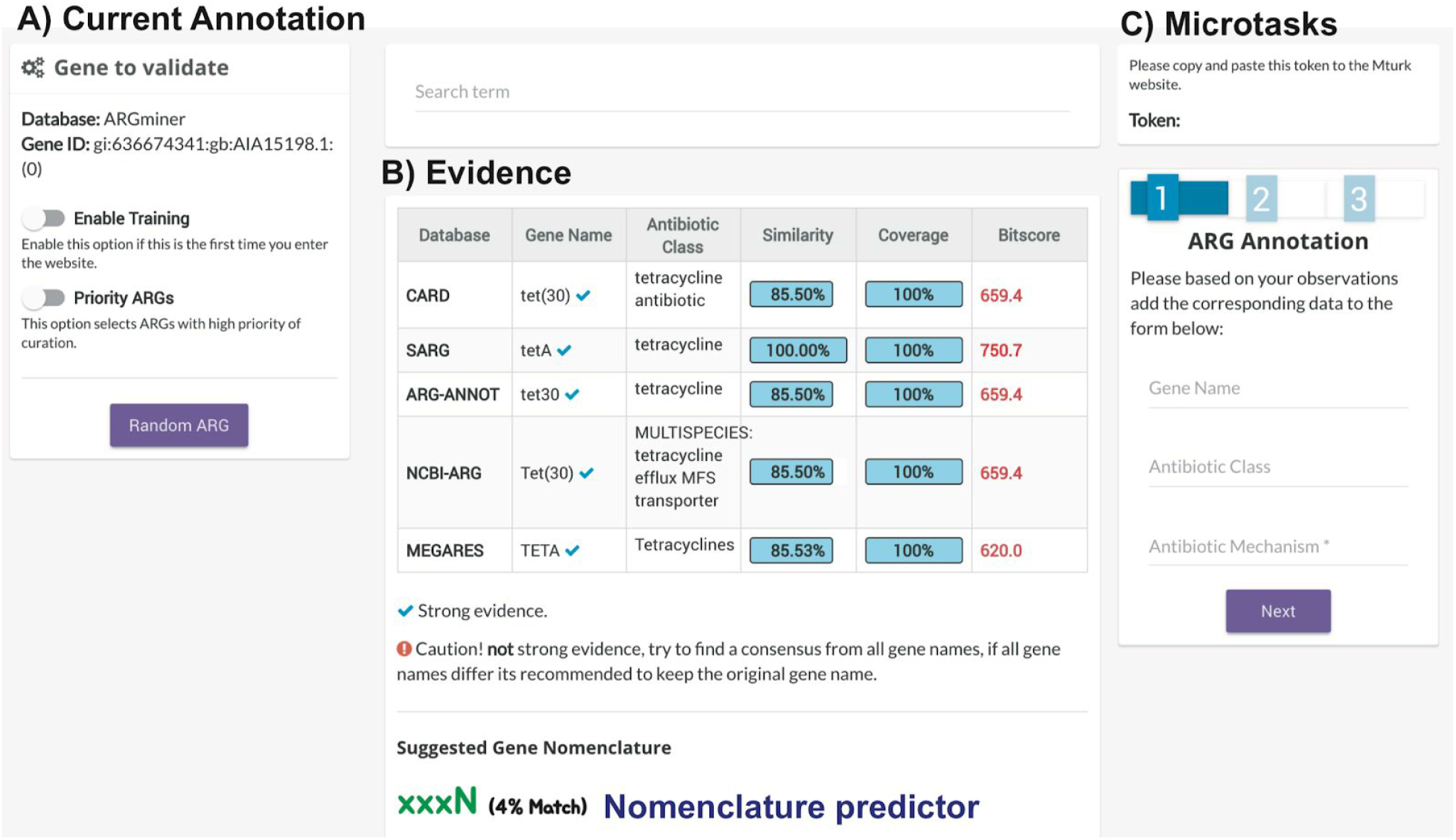
General overview of the ARG-miner platform. **A)** Current annotation. This panel contains the current information available for the ARG entry that requires validation. The “priority ARGs” option enables to curate ARGs in the database that have conflicting annotations. **B)** Evidence. This is the main panel and provides all of the metadata and information extracted from the different databases and resources. Note that in this panel there are colors that describe the relevance of each scoring metric. This is useful for users that are not familiar with alignment scores. **C)** Microtasks. This section contains the three microtasks needed for the ARG curation. It also contains real-time validation, which prompts error messages if the user inputs errors.

The web interface provides a training step for new users that is mandatory for AMT curators (required for getting a monetary reward). The goal of this step is to familiarize the curators with the platform environment by performing two microtasks. ARGminer also provides a list of problematic ARGs that have inconsistent annotations. These problematic ARGs are identified by comparing the annotation of the genes with their best hits from ARDB, CARD, and MEGARes. All tests performed during validation were completed using these problematic ARGs.

ARGminer also provides an administrative interface to update the ARG database. This interface comprises a set of figures that show the distribution of different labels as well as the MGE and pathogenic evidence scores. In this interface, ARGminer administrators are able to accept/reject the annotations made by the crowd and update the ARG database (**see Supplementary Figure 6**).

### Trust validation filter

Because of the unsupervised nature of crowdsourcing, users can provide erroneous feedback or just ignore the evidence and enter random inputs. Under an uncontrolled scenario, spammers can even get a monetary reward. More critically, too much random and/or erroneous feedback can increase the variance in ARG annotations and propagate annotation error. To circumvent this problem, ARGminer implements a trust-validation filter (see **Supplementary Material**) to evaluate whether the input corresponds to actual evidence or not. This score is computed in real time, and unless the user provides valid information, the system will not proceed to the next stage. **Supplementary Figure 7** shows an example of a user providing erroneous input for the antibiotic class field.

### Annotation score

ARGminer scores the curation of ARGs based on the majority voting strategy described in {Prill, 2011 #3090} weighted by the validation-filter score and the curator’s estimated evidence score and expertise (**Supplementary Material** for details). The majority voting strategy assigns to the ARG under investigation the most common label provided by the curators. In other words, labels are assigned based on the highest count or frequency, adjusted by the evidence, expertise, and trust-validation scores.

## RESULTS AND DISCUSSION

### Nomenclature prediction

A machine learning model to assist annotators to identify the right ARG nomenclature was embedded into the ARGminer platform to assists curators to determine the gene name nomenclature. To train the model, 17 different gene name shapes with at least 10 genes were identified (**see Supplementary Table 1**). Thereafter, to assess the performance of the nomenclature prediction module in ARGminer, precision and recall over the validation set were computed. Precision is the ratio of the number of correctly predicted gene names over the total number of genes in the validation data set, while recall is the ratio of the number of correctly predicted gene names over the total number of genes with this name. The model achieved a high precision of 0.87 ± 0.02 and recall of 0.87 ± 0.02 in the validation set composed of 452 entries that were not used during the training process. **Supplementary Table 2** shows examples of predicted gene names along with their input. For instance, the uniprot entry D3JX00 has been reported by different resources as aadA2, ant2ia, ANT3-DPRIME, and AadA2, the system recommends the gene name to have the shape xxxXN with a 63% match.

### Crowdsourcing curators

To assess the effectiveness of the crowdsourcing approach for ARG annotation, we evaluated three groups of curators with the following attributes:

1. A set of crowdsourcing curators from MTurk, referred to as AMT-Free. In this scenario, curators were paid $0.10 for each annotation, with the trust validation filter disabled to examine the reliability of the general crowd. Therefore, curators could input anything as feedback without restriction. A total of 100 annotations were requested from MTurk for this test.
2. A second batch of crowdsourcing curators from MTurk, referred to as AMT-Val. In this case the trust validation filter was enabled. The main purpose of this experimental group was to measure the effectiveness of the trust validation filter. In this scenario, a total of 200 annotations were requested from MTurk.
3. A group of users with general microbiological knowledge, with varying levels of experience in ARG research, referred to as **LAB**. This group consisted of Masters and Ph.D. students from a microbiology class at Virginia Tech. They completed this work as an assignment and did not receive any monetary reward. Here the annotations were performed with the validation filter on and each curator was requested to perform 15 annotations (540 microtasks in total). The goal of the LAB scenario was to compare its performance against the non-expert community of MTurk (AMT-Val, AMT-Free).

### Effectiveness of the trust validation filter

Spammers are curators that intend to obtain monetary reward by submitting invalid information, which is a major confounder to effective crowdsourcing. In the present study, although the ARGminer web site provides curators with detailed instructions about how to handle the annotation process, many of the **AMT-Free** curators submitted misleading and/or unrelated feedback. For the antibiotic category annotation task, curators must choose the antibiotic class to which they believe the gene belongs from a dropdown menu that contains a list of antibiotic classes. Results indicated that many **AMT-Free** curators simply selected the first option on the dropdown menu (aminoglycosides class), most likely without reading the evidence section of the web page. Thus, it was observed that most of the antibiotic class annotations under the AMT-Free group were labeled as aminoglycosides. This is a serious hurdle to accurate database curation and indicates the need for a real time control that guarantees accuracy of the annotation. In terms of performance, as expected, the AMT-Free group achieved very low scores for all annotations (**Figure 2**). However, not all curators were spammers. It was observed that few curators who performed more than ten microtasks also responded correctly and consistently to their observations and evidence. After integration of the trust validation filter, MTurk curators were monitored on their feedback (**AMT-Val** group) (see **Figure 2**). As a result, the performance of the AMT-Val curators improved significantly for all fields (antibiotic class, ARG name, and ARG mechanism) when compared to the **AMT-Free** group (P=1e-10 for annotation score distributions). Under the new restriction, MTurk curators were not allowed to continue with the microtask until their annotation was valid, as shown in **supplementary Figure 7**. Under this test, all nonsense input was eliminated and all annotations from the **AMT-Val** group corresponded to actual ARG evidence prompted by the web site. These results demonstrate the effectiveness of the trust validation filter for the control of spam annotations. In addition, it was imperative to test the performance of the MTurk curators against domain knowledge users. The main goal of this test case was to investigate whether a nonexpert crowd community (**AMT-Free** and **AMT-Val**) can perform a complex task with comparable outcomes as those of a group of curators with domain-knowledge (**LAB**). As expected, the LAB curators achieved a much higher average score (0.146) than the AMT-Free curators (0.06), but, surprisingly, a rather similar score to the AMT-Val curators was observed (0.114). This demonstrates that crowdsourcing is indeed a powerful alternative to manual inspection and annotation of ARGs by domain experts. As expected, MTurk annotations (AMT-Val) were characterized by a higher variance compared to the LAB group in all annotation fields, but the two distributions were not significantly different (Kolmogrov-Smirnov test: P > 0.05).

**Figure 2:**
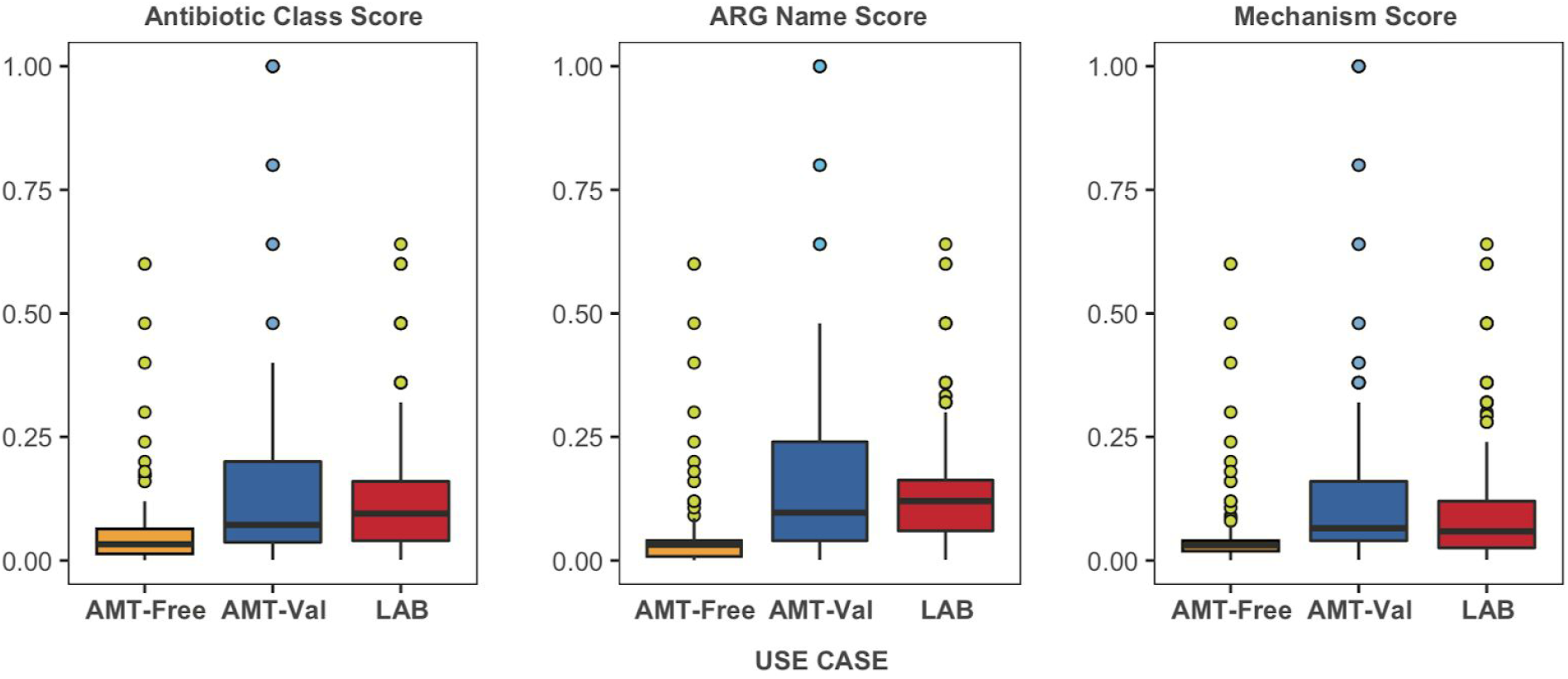
Annotation score of the three crowdsourced use cases (AMT-Free: Amazon MTurk curators without the true validation filter, AMT-Val: Amazon MTurk curators with the validation filter enabled and LAB: a group of curators with general microbiology domain knowledge and some antibiotic resistance knowledge. AMT-Val displayed the highest variance. However, this distribution was closer to that obtained by the curators with domain knowledge. Scores from the AMT-Free curators were the lowest among the three scenarios, indicating the ineffectiveness of the crowdsourcing annotation when the curator’s input was not validated.

### Effectiveness of the scoring strategy

To evaluate the quality of the scoring strategy, four genes were selected among the total set of curated genes and examined in greater detail, as illustrated in **Figure 3**. For instance, the UniProt entry A0A0D0NPG2 is a bifunctional polymyxin resistance protein, ArnA, involved in several biological processes including coenzyme binding, UDP-glucuronic acid dehydrogenase activity, lipid A biosynthetic process, and response to antibiotic. This protein builds the UDP-L-4-formamido-arabinose attached to lipid A, which is required for conferring resistance to polymyxin and cationic antimicrobial peptides {Huja, 2015 #3091}. From the crowdsourcing classification, both peptide and polymyxin antibiotic classes were identified, where polymyxin was characterized by a slightly higher score (**Figure 3A**). A closer look at the evidence from the antibiotic resistance databases (CARD, ARDB, and MEGARes) reveals a consensus of the gene towards the polymyxin antibiotic class. The evidence from the antibiotic resistance databases strongly suggests that the gene entry A0A127SI91 corresponds to a bl1-EC beta lactamase gene. **Figure 3C** illustrates different crowd classifications (including all evaluation scenarios). Note that beta-lactam is the dominant class with the highest annotation score. However, as a consequence of disabling the trust validation filter, several unrelated categories were accepted, such as aminoglycoside, MLS, multidrug, nitrofurantoin, polyamine, polymyxin, and even the word “yes”. One particularly interesting observation is the remarkable similarity among valid annotations. For instance, in **Figure 3D**, the gene AAC76733.1 was correctly assigned to multidrug as its best classification and to the “multi-drug resistance” category as its second best classification. Such small semantic differences are not detected by the trust validation filter. Therefore, under the validation interface, the administrators of ARGminer have the ability to validate or reject the annotations if needed. **Figure 3B** shows that most curators assigned the gene A0A0Q9QYU5 to the beta lactamase category. Note that the suggested name “beta_lactam” is the highest scored among all choices.

**Figure 3:**
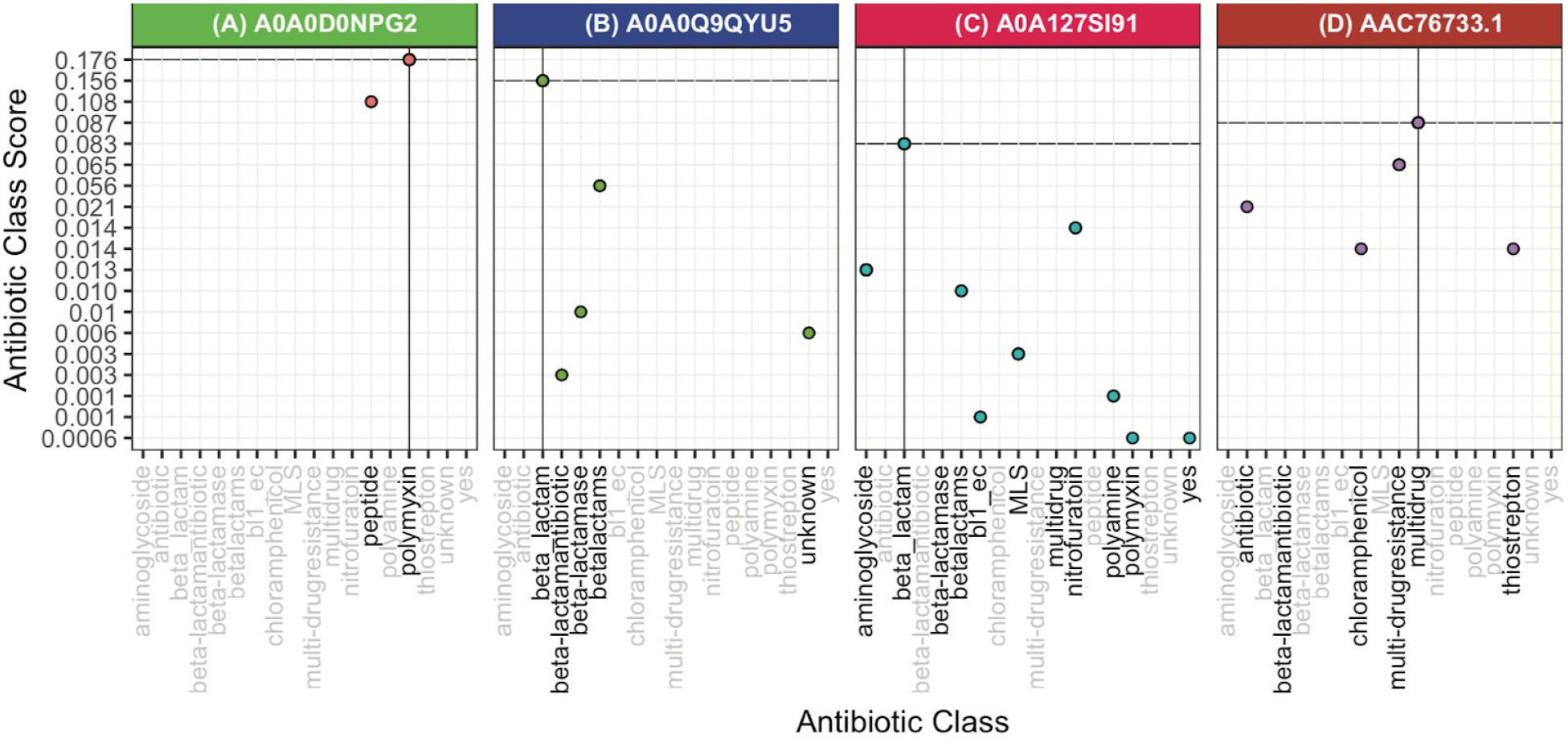
Distribution of the antibiotic class annotation by the crowdsourcing curators using the annotation score. X axis corresponds to the antibiotic resistance categories, where black labels indicate the categories reported by the curators and the top of each box corresponds to the ARG identifier.

**Figure 4** shows the crowdsourced score for the ARG name classification. As seen for the antibiotic category annotation, there are cases where the annotations are semantically highly similar. For instance, the gene A0A127SI91 was tagged as bl1_ec, bl1-ec, or blaec, all corresponding to the bla1-EC gene name (**Figure 4C**). Note that all these labels were ranked higher than the other gene names (macb, baca, ba1) and all the unrelated tags such as “mm-58”, “15”, “yes”, and “middle”. Also, all unrelated annotations were ranked low by the scoring strategy.

**Figure 4:**
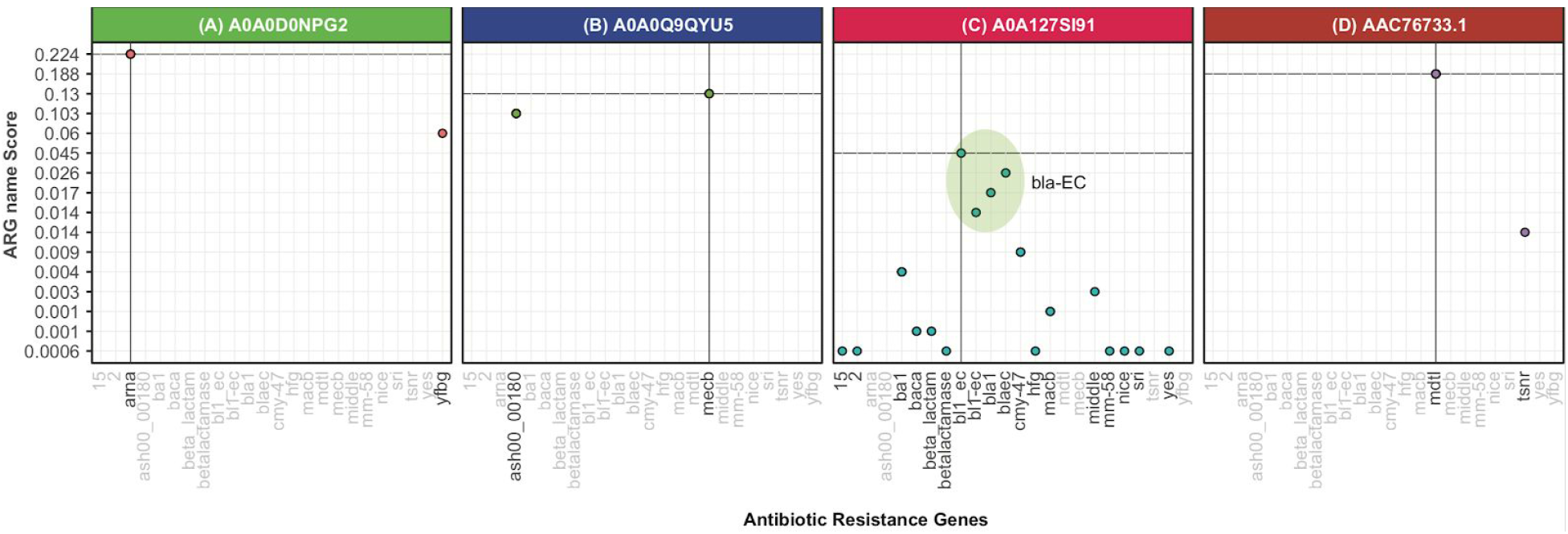
Distribution of the prediction of ARG names. ARG names are represented on the x axis and the y axis indicates the corresponding annotation score. The top of each box corresponds to the ARG identifier.

Although identification of the antibiotic category for the gene A0A0Q9QYU5 was straightforward, the detection of its gene name is challenging, primarily because the metadata of this entry does not include the gene name and because the identity of its best hit alignments is below 30%. This indicates that the gene has a potential homology to known ARGs. Two ARG databases (CARD and MEGARes) show a significant best hit e-value (<1e-22) over the mecB gene. For this example, 50% of the curators annotated the gene as mecB whereas the other 50% annotated it as ash00_000180. Also, curators yielded a higher confidence for the *mecB* gene (2.6 average confidence score) compared to the ash00_000180 (2.3 average confidence score). As a result, mecB achieved a slightly higher score. To document any uncertainty, ARGminer recommends that users retain the original label if the evidence is not convincing. For the other examples (**Figure 4A and 4D**), the crowd classified the gene names according to the observed evidence.

### Annotation analysis

To assess the accuracy of the crowdsourcing annotation, genes that were inspected by fewer than 10 curators were removed from the total pool of classified genes. Then, a total of 35 genes were identified and manually curated by three domain experts according to antibiotic class and gene name. It was found that experts achieved an annotation pairwise correlation of 0.96± 0.02, indicative of an almost perfect classification agreement. Genes that were classified to the same label by at least two experts were used as the gold standard data set (**see Supplementary Table 3**). This benchmark was then used to measure the performance of the crowdsourcing curators where labels were selected based on the greatest annotation score.

The crowdsourcing classification of the antibiotic classes was essentially just as accurate as the expert annotation (94% Positive Predictive Value - PPV). In other words, 33 out of 35 genes labeled via crowdsourcing matched the expert classification (see Supplementary File 1). The genes for which the curators failed to identify the correct antibiotic class were a quinolone ARG annotated as multidrug (YP_001693238) and a multidrug gene annotated as quinolone (NP_358469.1). The classification of the ARG names proved to be a challenging task. Indeed, experts did not fully agree about the correct name of five ARGs (**see Supplementary Table 3**). However, only one of those conflicting genes was assigned a different classification assigned by all three experts. This gene corresponded to a macrolide gene (AFU35065.1), which was tagged as Isa, Isa-A, and Isa-E by the three experts. Thus, this gene entry was removed for the gene name analysis comparison and the final control data set contained 34 genes. When comparing the gene name annotation from the crowdsourcing curators, their prediction had a 97% PPV (**see Supplementary File 2**). This indicates that only one gene was not correctly classified by the crowd (J2LT98). By examining the details of this gene in ARGminer, all three ARG databases agreed that the gene belonged to the SHV group, with markedly high scores. However, CARD labeled it as the SHV variant 1 (SHV-1), ARDB labeled it as variant 2 (SHV-2), and MEGARes labeled it as the SHV group, without specification of a variant. An interesting aspect with respect to this particular ARG is that variants are defined by specific amino acid modifications {Paterson, 2003 #3093}, thus these genes are highly similar and identifying the correct variant by using sequence alignment is a particularly difficult task, as shown in **Supplementary Figure 8**. This aspect has the potential to confuse curators when classifying genes that are highly similar. Interestingly, by examining the crowd results, curators were able to discard the SHV variant 2 (99.3% identity), but they were not able to differentiate between the SHV variant 1 and the SHV group (both have the same score). These results suggest that crowdsourcing curators are able to follow the correct track, even in the face of complex tasks. Interestingly, the gene name prediction model assigned the gene name nomenclature to XXX-N with a probability of 0.79, which corresponds to the nomenclature followed to name SHV beta-lactamase genes. Because of the risk of propagation errors, the updating process of ARGs requires administrator approval for new database releases in the ARGminer web site. Overall, the crowd exhibited performance comparable with the experts, but in much less time. These results suggest that crowdsourcing annotation is a strong alternative to manual classification and validation of ARGs by domain experts.

### Expertise and confidence

ARGminer asks users to rate their own expertise in the analysis of ARGs on a scale of 0 to 5. **Figure 5A** shows the distribution of the expertise score against the correct or incorrect annotations for the antibiotic category classification (including all scenarios: AMT-Free, AMT-VAL, and LAB). Surprisingly, it is clear that having expert knowledge does not really make a difference in the quality of the classification. Indeed, because of the open nature of AMT, most of the curators are not experts and have little knowledge about ARGs. From **Figure 6A**, it is also evident that the proportion of correct annotations was higher compared to the incorrect classifications (the size of the dot indicates the number of annotations). This result suggests that accurate detection of the correct antibiotic resistance category does not necessarily require domain expertise. On the other hand, curators were also required to rate their confidence in the annotation. **Figure 5B** shows that self-rated confidence is a strong predictor of the quality of the annotation. The distribution of the confidence score indicates that higher confidence correlates with more accurate results. For instance, 95% of the curators who rated their confidence with 5 stars achieved correct annotation and only 5% missed. This strongly suggests that the confidence score is a superior indicator of correct annotation over the expertise score.

**Figure 5:**
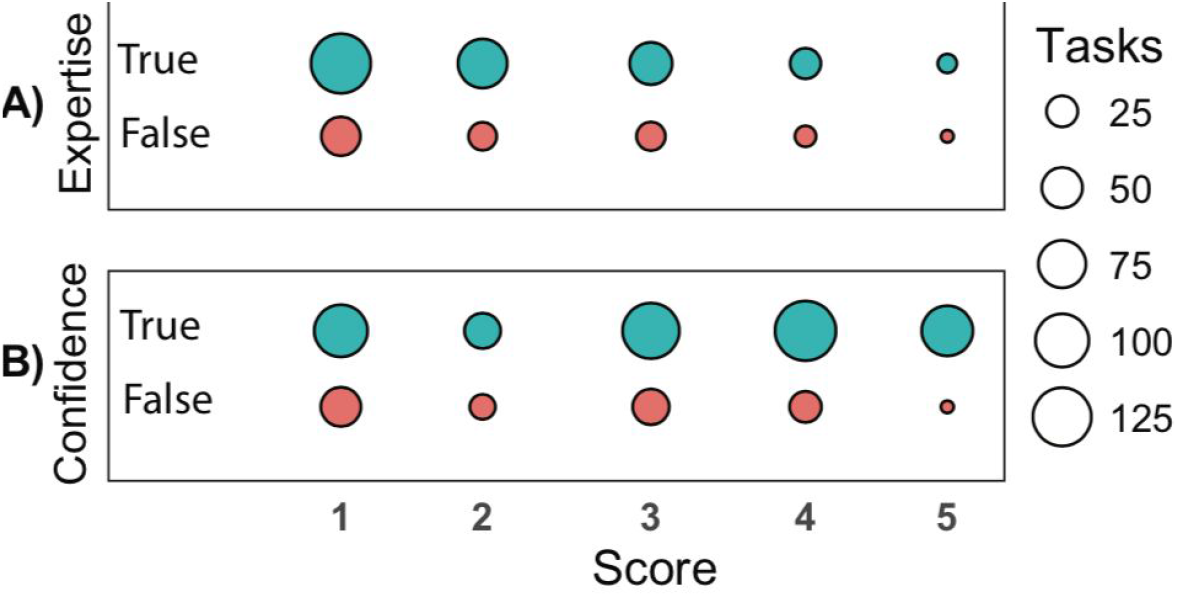
Expertise and confidence levels of the curators. The size of the points indicates the number of tasks; the x axis corresponds to the score level and the y label shows the expertise and confidence parameters. Color depicts correct and incorrect classifications.

## CONCLUSIONS

Here we develop, launch, and validate a new web platform, ARGminer, a powerful system that harnesses the power of crowdsourcing for advancing robust and comprehensive curation of ARGs. ARGminer enables easy access to key relevant information pertaining to ARGs, including metadata, evidence of ARGs being carried by pathogens, and the possibility of ARGs being mobilized by MGEs. Further, it enables a simple, but powerful, tool for the curation of ARGs designed to provide accurate information represented in a noncomplex way that can be validated by users without the requirement of domain knowledge. Results demonstrated not only that crowdsourcing curators yield curations as accurate as experts, but they are also more efficient than ARG-domain experts. Thus, ARGminer opens the possibility of a truly comprehensive, accurate, and perpetually up-to-date publicly-available ARG database.

**Supplementary Material File 1:**

Antibiotic classification of all gene entries from the expert validated dataset.

**Supplementary Material File 2:** Antibiotic resistance names annotation of all gene entries from the expert validated dataset.

