## Supplementary material for "ARGminer: A web platform for crowdsourcing-based curation of antibiotic resistance genes"

**Annotation Score Strategy**

ARGminer scores potential ARG curations based on the majority voting strategy ($M$) described in {Prill, 2011 #3090} weighted by the trust validation-filter score ($V$), worker's confidence ($C$) and expertise ($E$). To describe these values, let us assume we want to score the names of a gene $g_i$, that has been inspected by $W_n$ crowdsourcing workers who had available a total of $L$ different gene names to choose from.

The majority voting strategy, the expertise, the confidence and the trust validation scores can be arranged in a matrix where columns represent the crowdsourcing workers and rows represent each label (See Supplementary **Figure 1**). Thus, to measure the score for each labels, we need to sum up the row scores and divide them by the total number of workers. The final annotation score for the label $L_i$ is equal to the multiplication of the individual scores ($A_i = M_i * E_i * C_i * V_i$).

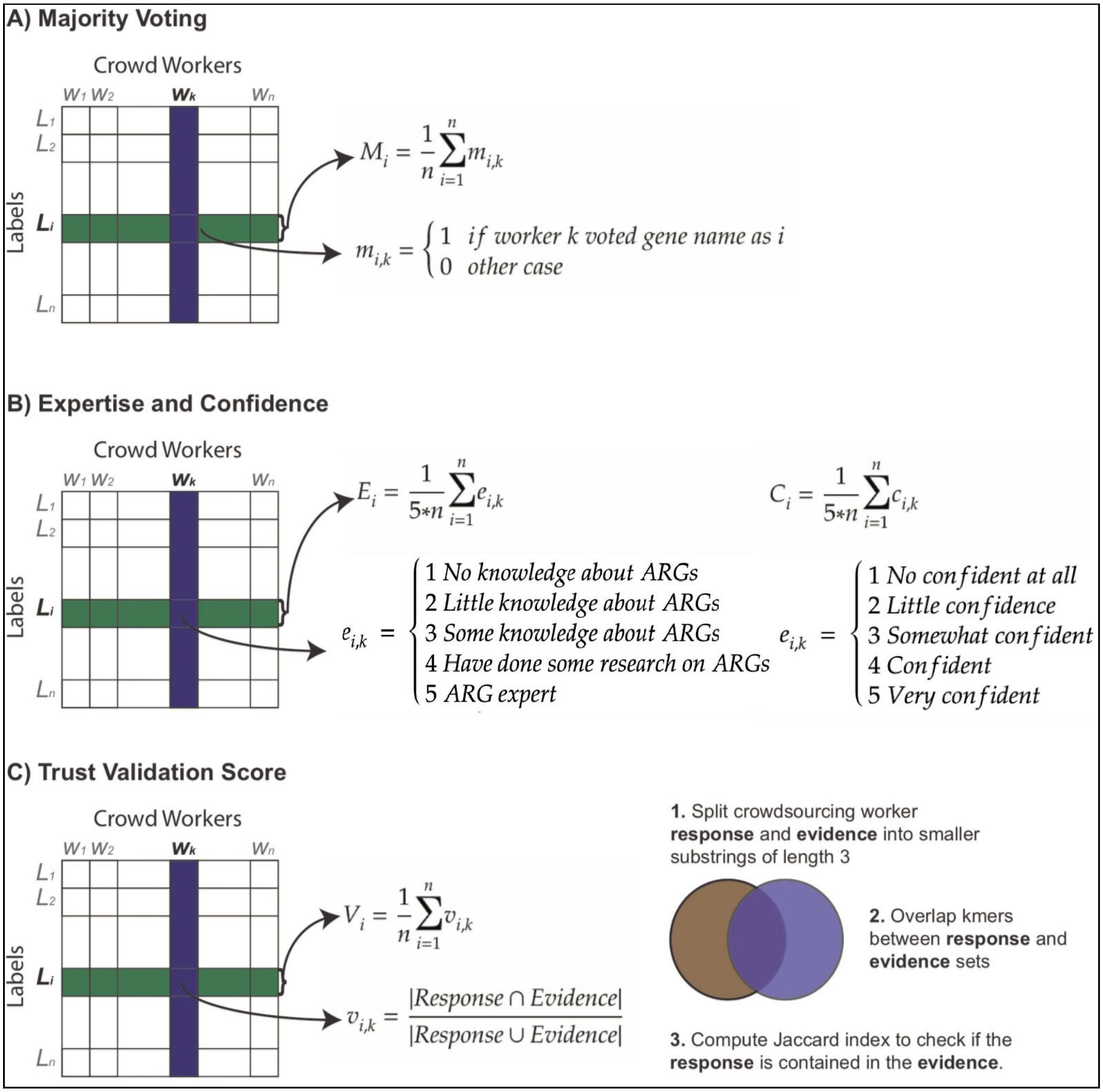


**Supplementary Figure 1**: Scoring strategy used for annotation of ARGs. **A)** Majority voting score obtained from the total number of workers. **B)** Expertise and confidence normalized from 0 to 1 scores. **C)** Strategy to compute the score for the trust validation-filter.

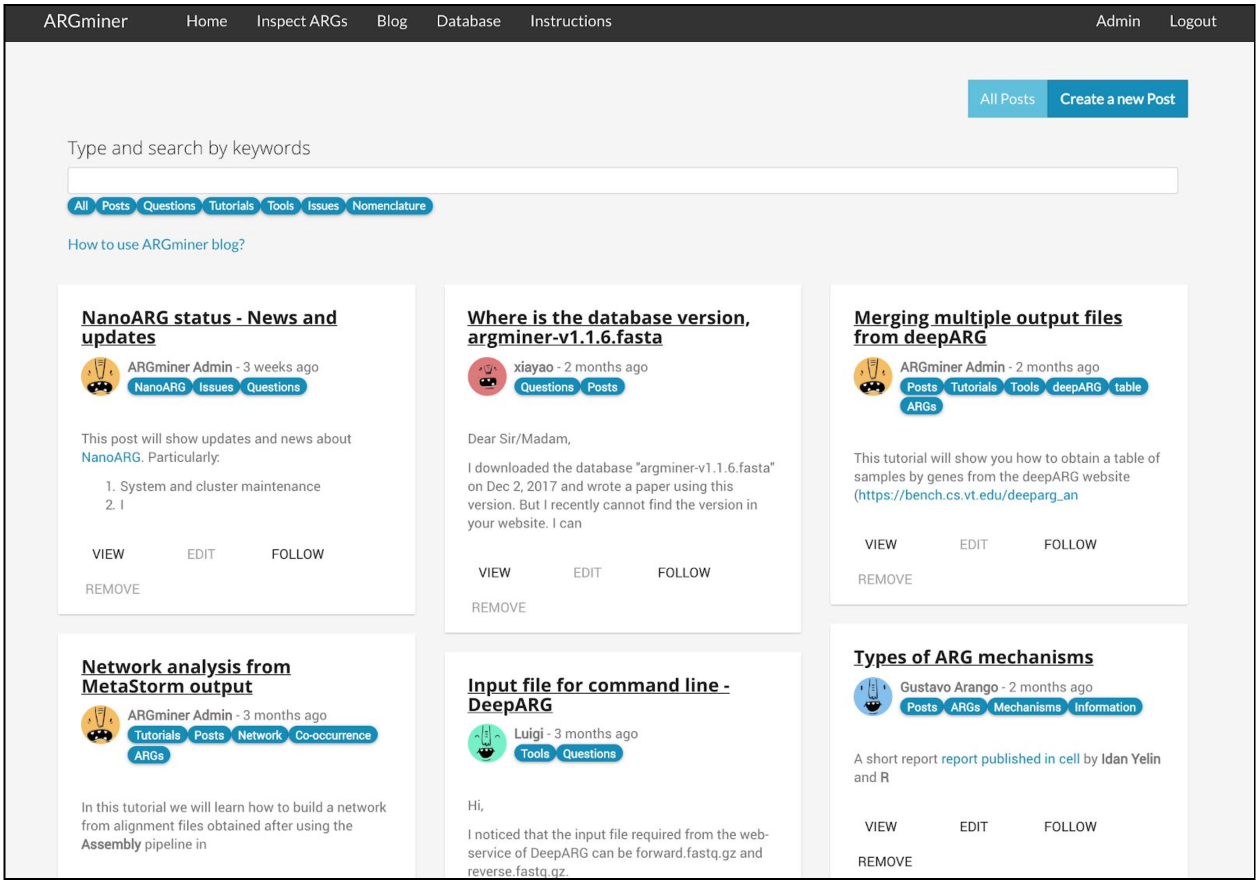


**Supplementary Figure 2**: ARGminer blog available for users to upload questions, posts, tutorials about analysis of ARGs or general nomenclature questions.

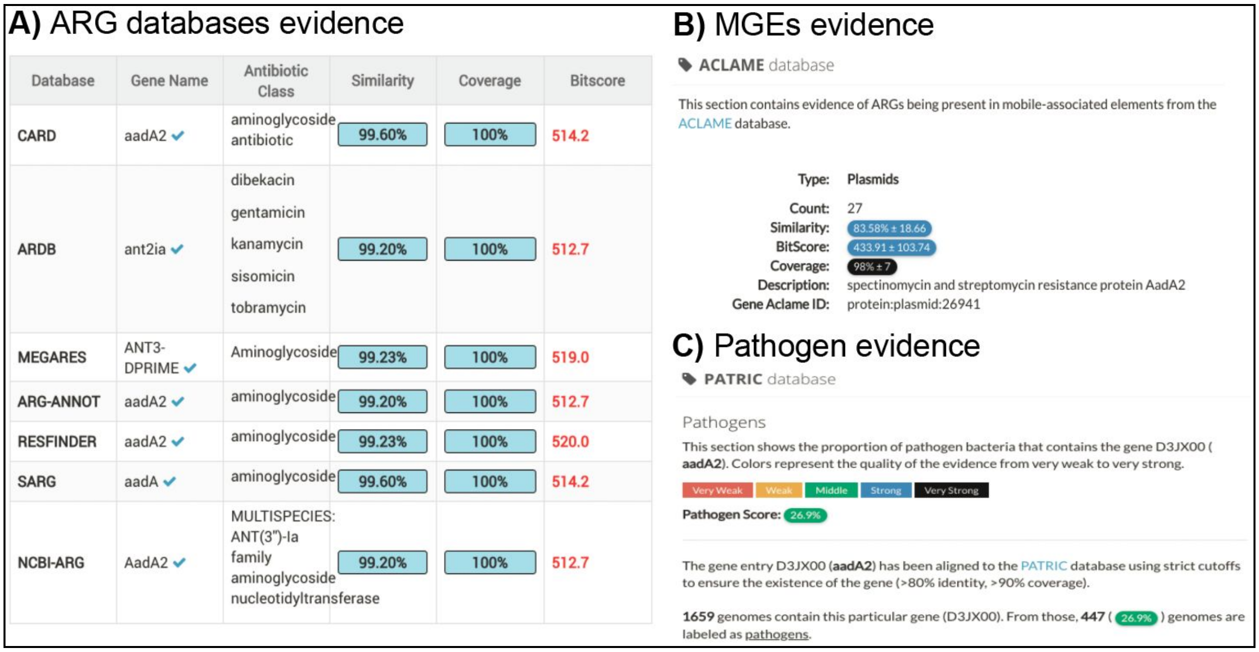


**Supplementary Figure 3**: Evidence of ARGs in ARGminer platform. **A)** Antibiotic resistance database hits. **B)** Mobile Genetic Elements. **C)** Evidence the ARG is being carried by a pathogen.

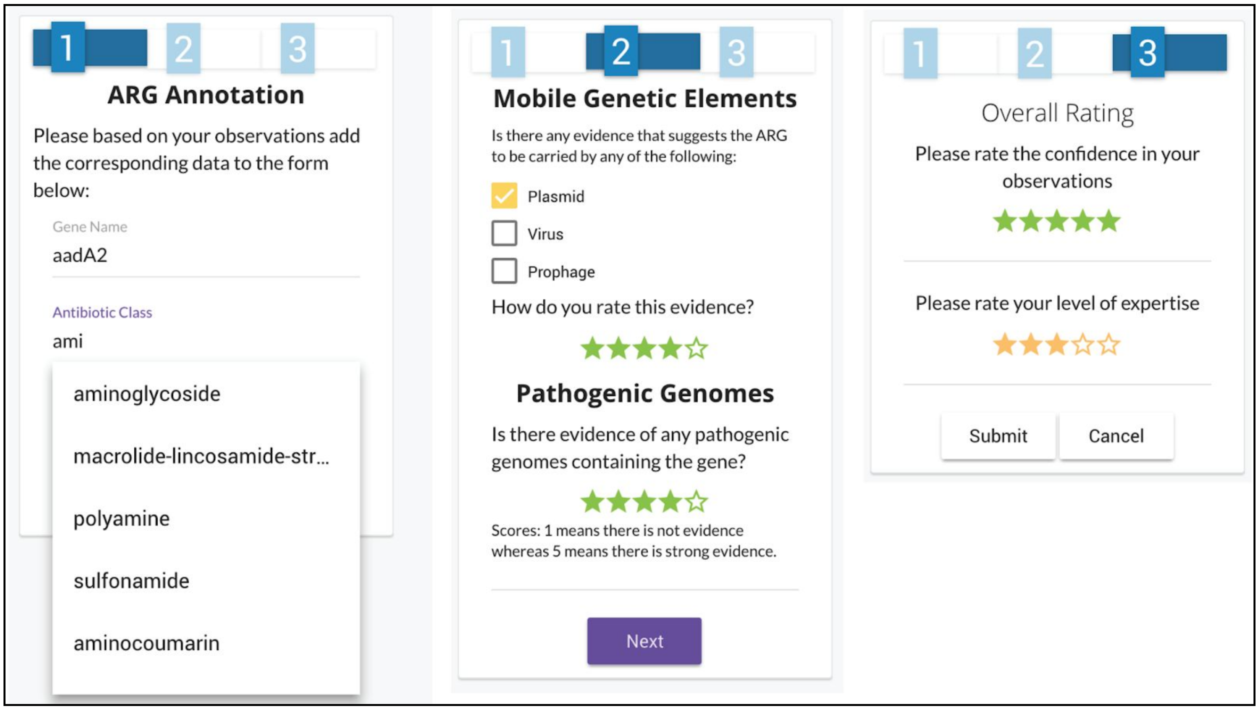


**Supplementary Figure 4**: Annotation process: First, ARG name, antibiotic class and ARG mechanism are requested to be filled up by worker. Then, workers are requested to check the evidence about MGEs and pathogens and score their observations. Finally, workers are requested to rate their confidence and expertise from 1 to 5.

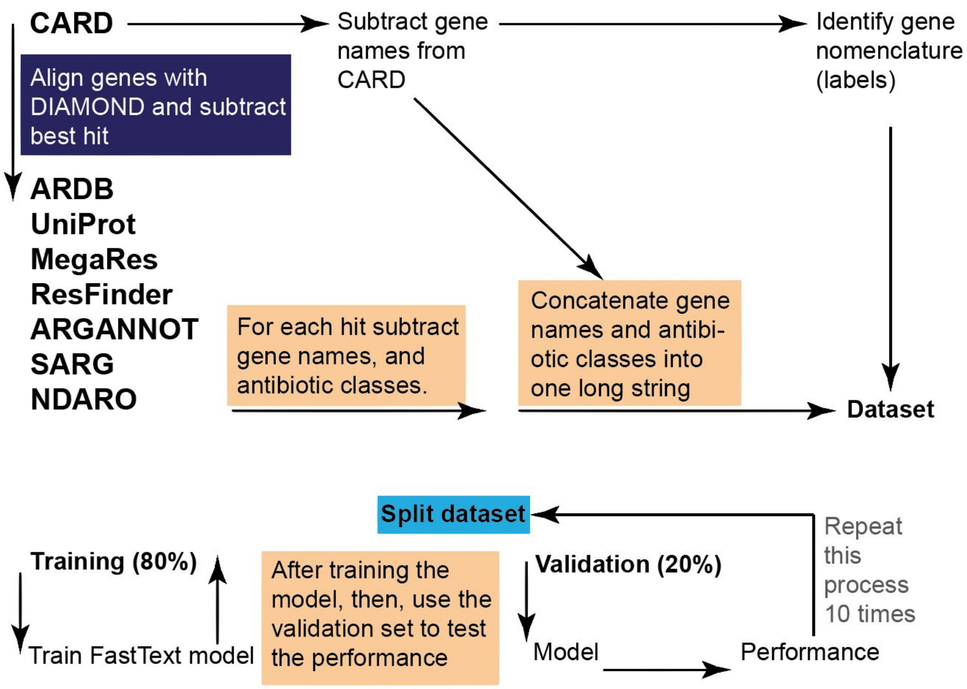


**Supplementary Figure 5**: General framework used for building the gene nomenclature dataset and to build the machine learning predictor using the natural language processing library FastText.

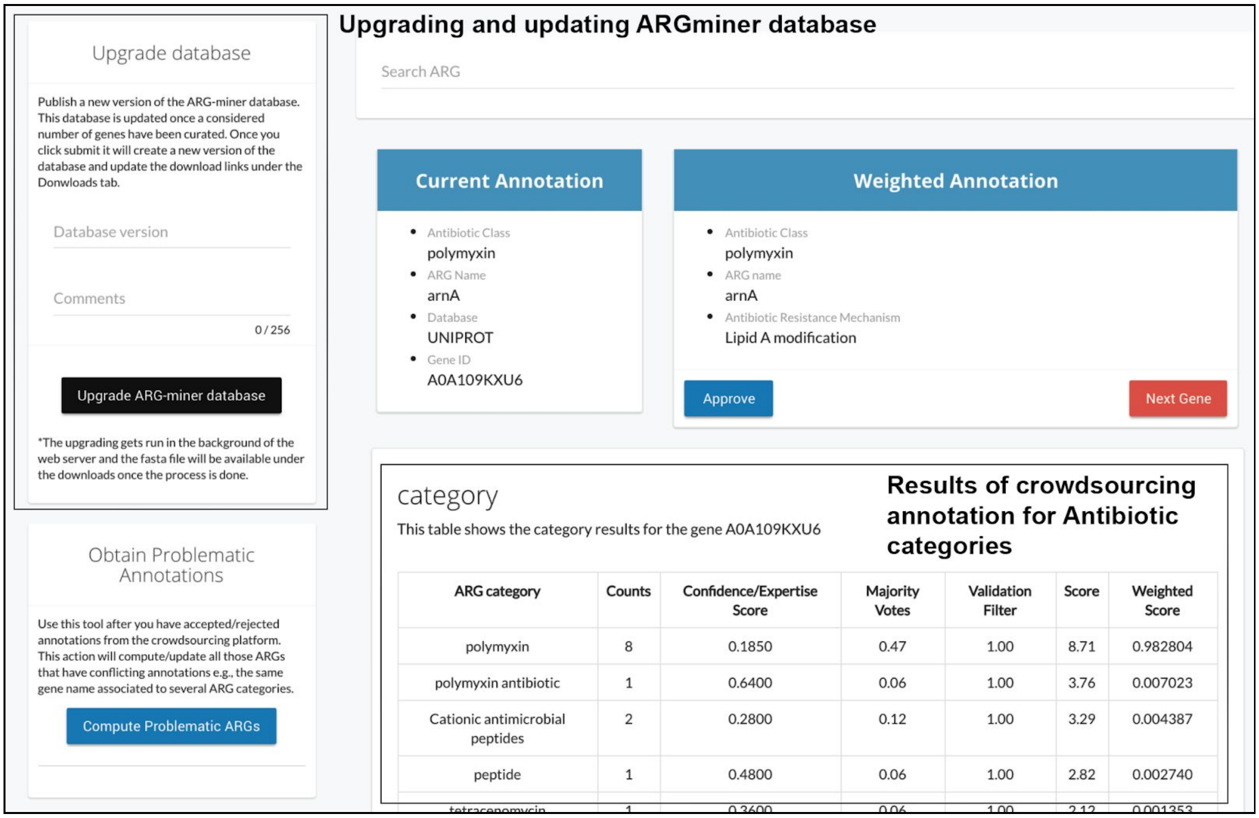


**Supplementary Figure 6**: Administration page of ARGminer. Once a gene is reviewed by a select number of workers, administrators of ARGminer can evaluate their annotations and approve or disapprove the crowd classification.

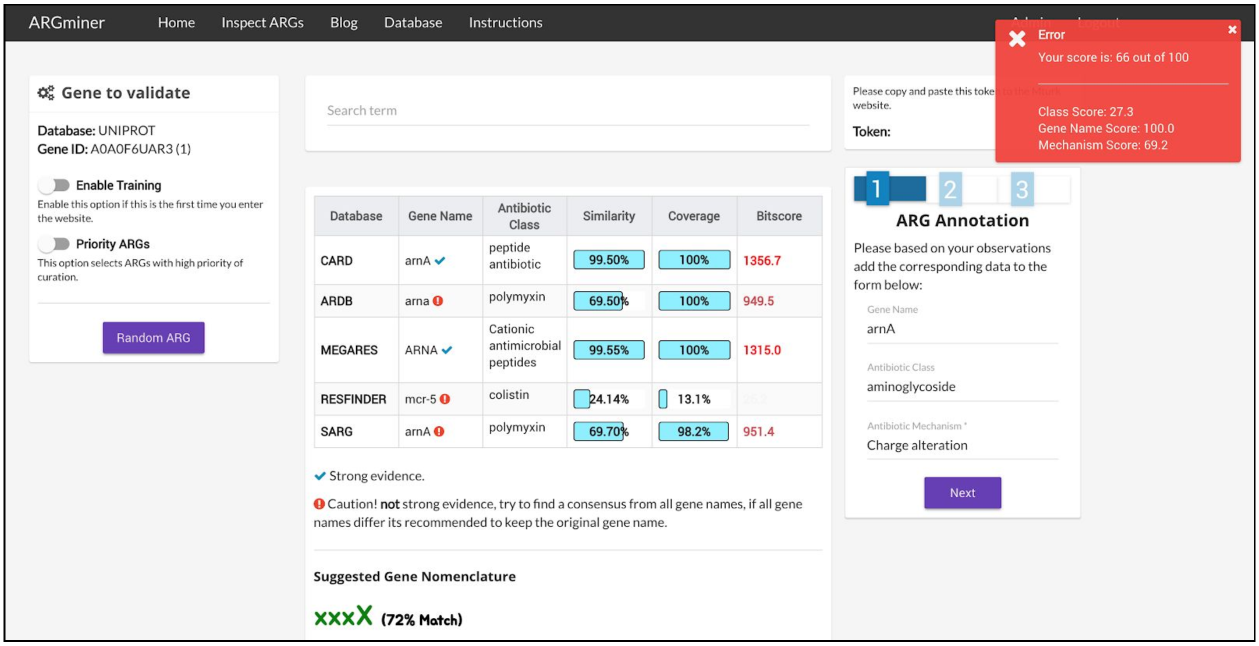


**Supplementary Figure 7**: Trust validation blocks entries with values that are not in the evidence section.

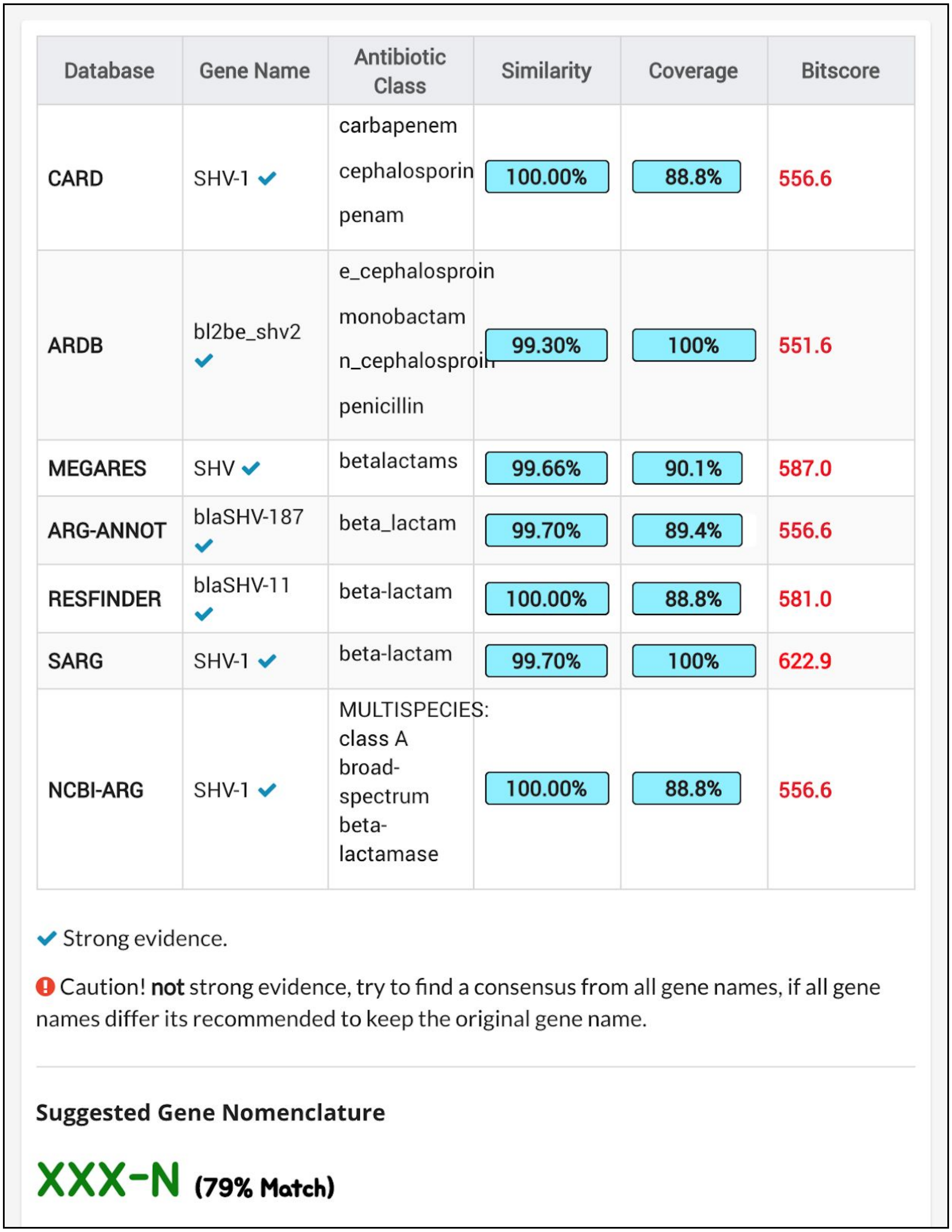

| Database | Gene Name | Antibiotic Class | Similarity | Coverage | Bitscore |
|---|---|---|---|---|---|
| CARD | SHV-1 ✔ | carbapenem cephalosporin penam | 100.00% | 88.8% | 556.6 |
| ARDB | bl2be_shv2 ✔ | e_cephalosproin monobactam n_cephalosproin penicillin | 99.30% | 100% | 551.6 |
| MEGARES | SHV ✔ | betalactams | 99.66% | 90.1% | 587.0 |
| ARG-ANNOT | blaSHV-187 ✔ | beta_lactam | 99.70% | 89.4% | 556.6 |
| RESFINDER | blaSHV-11 ✔ | beta-lactam | 100.00% | 88.8% | 581.0 |
| SARG | SHV-1 ✔ | beta-lactam | 99.70% | 100% | 622.9 |
| NCBI-ARG | SHV-1 ✔ | MULTISPECIES: class A broad-spectrum beta-lactamase | 100.00% | 88.8% | 556.6 |

✔ Strong evidence.

! Caution! **not** strong evidence, try to find a consensus from all gene names, if all gene names differ its recommended to keep the original gene name.

**Suggested Gene Nomenclature**

**XXX-N (79% Match)**

**Supplementary Figure 8**: ARGminer Uniprot gene entry J2LT98 is recognized by major databases. The nomenclature predictor suggests the name should be written as SHV-N where N depicts the gene variant for the SHV beta lactamase family.

| Nomenclature | ARG counts | ARG Example |
|---|---|---|
| XXX-N | 1153 | CMY-2 |
| xxxX | 338 | ileS |
| XXX-X-N | 174 | CTX-M-124 |
| XxxXN | 102 | FosA4 |
| XxxX | 86 | OqxA |
| xxxXN | 65 | dfrB6 |
| xxxXX | 53 | vanXA |
| XXX(N')-Xx | 37 | AAC(6')-Ip |
| xxx | 29 | cat |
| XXXX-N | 25 | HERA-1 |
| xxxN | 24 | tet32 |
| XXX-N-N | 23 | OXY-1-6 |
| XXX(N)-Xx | 17 | ANT(6)-Ia |
| xxxx | 16 | mdtl |
| Xxx(N) | 15 | Erm(35) |
| XXX(N')-XXx | 12 | AAC(6')-IIa |
| xxx-N | 11 | arr-8 |

**Supplementary Table 1**: Nomenclature shapes detected in CARD. The table shows shapes that has at least 10 genes.

| Training text | Label |
|---|---|
| CTX-M beta-lactamase cephalosporin antibiotic inactivation bl2be_ctxm ceftazidime cephalosporin_i cephalosporin_ii cephalosporin_iii monobactam penicillin betalactams CTX beta_lactam blaCTX-M-15 beta-lactam blaCTX-M-142 beta-lactam CTX-M-142 class A extended-spectrum beta-lactamase CTX-M-142 | XXX-X-N |
| vanY glycopeptide resistance gene cluster glycopeptide antibiotic antibiotic target alteration vanyg vancomycin Glycopeptides VANYG glycopeptide vanY-B glycopeptide VanXY_C2 vancomycin vanG MULTISPECIES: D-Ala-D-Ala carboxypeptidase VanY-G1 | xxxXXN |
| major facilitator superfamily (MFS) antibiotic efflux pump tetracycline tetracycline antibiotic efflux pump complex or subunit conferring antibiotic resistance antibiotic efflux tetc tetracycline Tetracyclines TETA tetracycline tet30 tetracycline tet(30) tetracycline tetA MULTISPECIES: tetracycline efflux MFS transporter Tet(30) | xxx(N) |

**Supplementary Table 2**: Examples of entries in the dataset for the nomenclature predictor. The training test consists of merging the information from different databases into a long string whereas the label corresponds to the gene name shape.

| Curator | Entry | Antibiotic Class | ARG |
|---|---|---|---|
| A | A0A0D0NPG2 | polymyxin | arnA |
| B | A0A0D0NPG2 | polymyxin | arnA |
| C | A0A0D0NPG2 | polymyxin | arnA |
| A | A0A0Q9QYU5 | beta lactam | mecB |
| C | A0A0Q9QYU5 | beta lactam | mecB |
| B | A0A0Q9QYU5 | beta lactam | mecB |
| A | A0A109KXU6 | polymyxin | arnA |
| B | A0A109KXU6 | polymyxin | arnA |
| C | A0A109KXU6 | polymyxin | arnA |
| C | A0A127SI91 | beta lactam | bl1_ec |
| B | A0A127SI91 | beta lactam | bl1_ec |
| A | A0A127SI91 | beta lactam | bl1_ec |
| A | AAB53445.1 | streptothricin | SAT-4 |
| B | AAB53445.1 | streptothricin | SAT-4 |
| C | AAB53445.1 | streptothricin | SAT-4 |
| A | AAC43969.1 | chloramphenicol | opcM |
| B | AAC43969.1 | multidrug | opcM |
| C | AAC43969.1 | multidrug | opcM |
| A | AAC60781.1 | cationic | rosA |
| B | AAC60781.1 | polymyxin | rosA |
| C | AAC60781.1 | polymyxin | rosA |
| A | AAC76733.1 | multidrug | mdtL |
| B | AAC76733.1 | multidrug | mdtL |
| C | AAC76733.1 | multidrug | mdtL |
| B | AAK51920 | aminoglycoside | aac(6')-Ib |
| C | AAK51920 | aminoglycoside | aac(6')-Ib |
| A | AAK51920 | aminoglycoside | aac(6')-Ib |
| A | AAM15533.1 | aminoglycoside | PvrR |

| | | | |
|---|---|---|---|
| B | AAM15533.1 | aminoglycoside | pvrR |
| C | AAM15533.1 | multidrug | pvrR |
| A | AAN80811 | multidrug | emrE |
| B | AAN80811 | multidrug | emrE |
| C | AAN80811 | multidrug | emrE |
| A | AAR84672.1 | glycopeptide | vanRC |
| B | AAR84672.1 | glycopeptide | vanRF |
| C | AAR84672.1 | glycopeptide | vanRF |
| A | ABA71733.1 | glycopeptide | vanTG |
| B | ABA71733.1 | glycopeptide | vanTG |
| C | ABA71733.1 | glycopeptide | vanTG |
| A | ABB43029.1 | aminoglycoside | AAC(3)-IV |
| C | ABB43029.1 | aminoglycoside | AAC(3)-IV |
| B | ABB43029.1 | glycopeptide | AAC(3)-IV |
| C | ABC54722 | aminoglycoside | aac(6')-Ib10 |
| B | ABC54722 | glycopeptide | aac(6')-Ib10 |
| A | ABC54722 | aminoglycoside | aac(6')-Ib10 |
| A | ABQ96629 | sulfonamide | sul1 |
| B | ABQ96629 | sulfonamide | sul1 |
| C | ABQ96629 | sulfonamide | sul1 |
| A | AFU35065.1 | MLS | lsa |
| B | AFU35065.1 | MLS | lsaA |
| C | AFU35065.1 | multidrug | lsaE |
| C | B5ULZ6 | beta lactam | bl2a_iii2 |
| B | B5ULZ6 | beta lactam | bl2a_iii2 |
| A | B5ULZ6 | beta lactam | bla1 |
| A | BAC11911.1 | multidrug | emeA |
| B | BAC11911.1 | multidrug | emeA |
| C | BAC11911.1 | multidrug | emeA |

| | | | |
|---|---|---|---|
| A | D3V1W5 | polymyxin | pmrF |
| B | D3V1W5 | polymyxin | pmrF |
| C | D3V1W5 | polymyxin | pmrF |
| C | F0JWD5 | polymyxin | anrA |
| A | F0JWD5 | polymyxin | arnA |
| B | F0JWD5 | polymyxin | arnA |
| A | J2LT98 | beta lactam | SHV |
| C | J2LT98 | beta lactam | shv-1 |
| B | J2LT98 | beta lactam | shv-1 |
| A | NP_251217.1 | multidrug | mdtB |
| B | NP_251217.1 | multidrug | muxB |
| C | NP_251217.1 | multidrug | muxB |
| A | NP_253677.1 | multidrug | emrE |
| B | NP_253677.1 | multidrug | emrE |
| C | NP_253677.1 | multidrug | emrE |
| B | NP_358469.1 | Acriflavin | pmrA |
| A | NP_358469.1 | multidrug | pmrA |
| C | NP_358469.1 | multidrug | pmrA |
| A | NP_415434.1 | multidrug | msbA |
| B | NP_415434.1 | multidrug | msbA |
| C | NP_415434.1 | multidrug | msbA |
| B | Q1RPS3 | aminoglycoside | aadA |
| C | Q1RPS3 | aminoglycoside | aadA |
| A | Q1RPS3 | aminoglycoside | ant |
| A | YP_001693238 | multidrug | oqxB |
| B | YP_001693238 | Quinolone | oqxB |
| C | YP_001693238 | Quinolone | oqxB |
| A | YP_002635521 | multidrug | mepA |
| B | YP_002635521 | multidrug | mepA |

| | | | |
|---|---|---|---|
| C | YP_002635521 | multidrug | mepA |
| B | YP_042788 | multidrug | bmr |
| C | YP_042788 | multidrug | bmr |
| A | YP_042788 | multidrug | norA |
| B | YP_490697.1 | aminoglycoside | acrD |
| C | YP_490697.1 | aminoglycoside | acrD |
| A | YP_490697.1 | multidrug | acrD |
| A | ZP_02377226 | multidrug | ceoA |
| B | ZP_02377226 | multidrug | ceoA |
| C | ZP_02377226 | multidrug | ceoA |
| A | ZP_02685589 | multidrug | mdfA |
| B | ZP_02685589 | multidrug | mdfA |
| C | ZP_02685589 | multidrug | mdfA |
| A | ZP_04003442 | multidrug | emrE |
| B | ZP_04003442 | multidrug | emrE |
| C | ZP_04003442 | multidrug | emrE |
| A | ZP_04679156 | multidrug | mepA |
| B | ZP_04679156 | multidrug | mepA |
| C | ZP_04679156 | multidrug | mepA |

**Supplementary Table 3**: Gold standard comparison. Classification of selected genes by curators A, B and C.
