## Supplementary figures and images for "ARGminer: A web platform for crowdsourcing-based curation of antibiotic resistance genes"

### SF 1

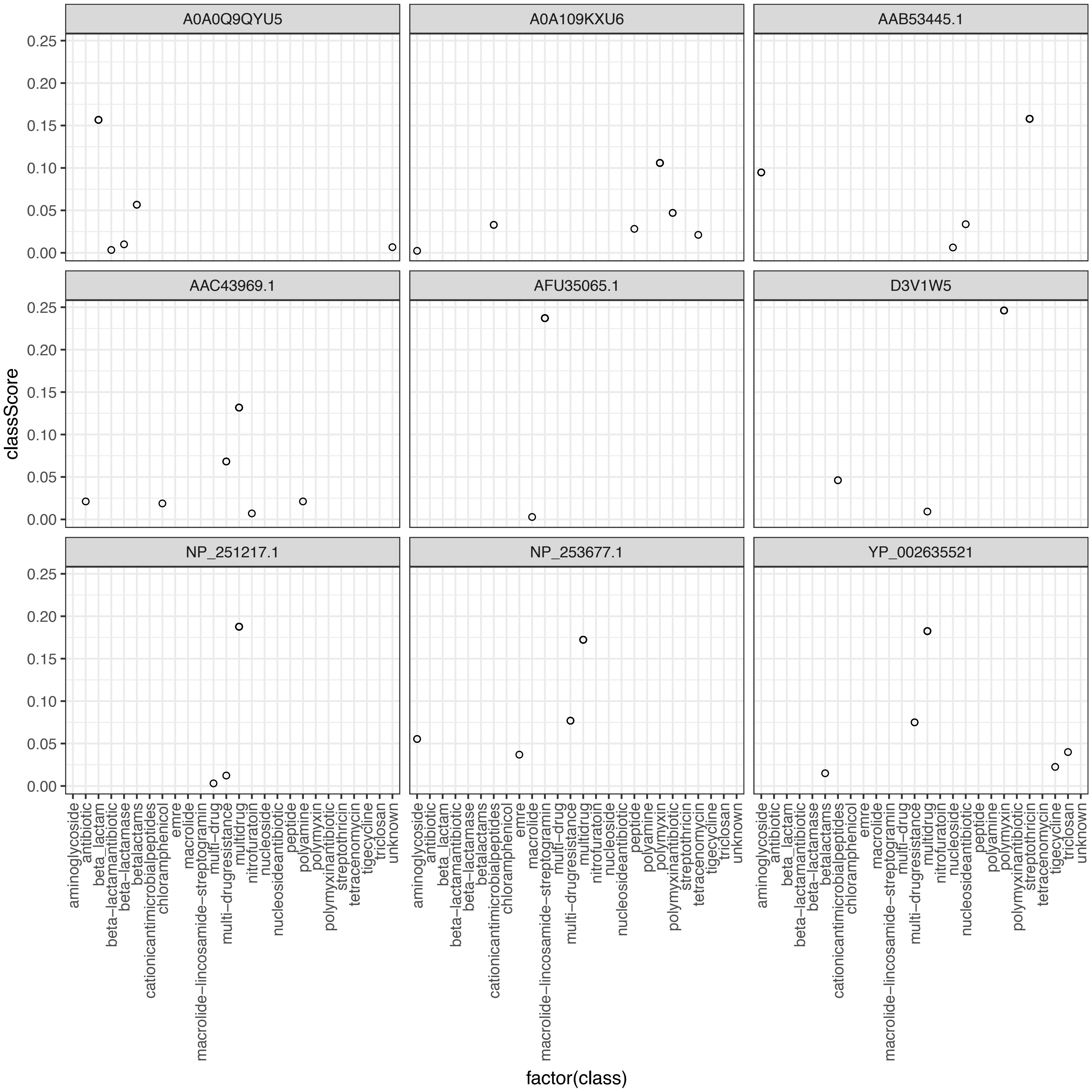

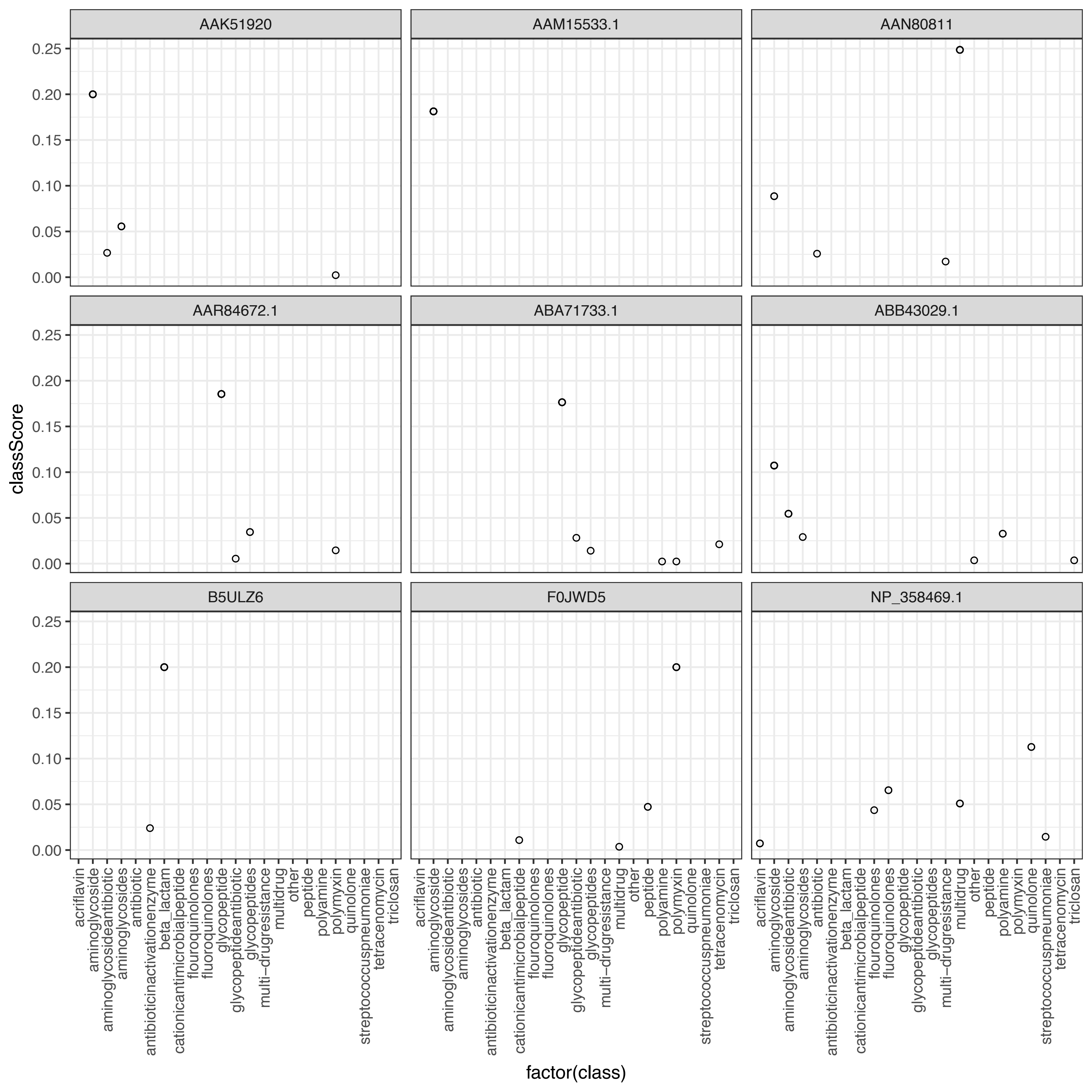

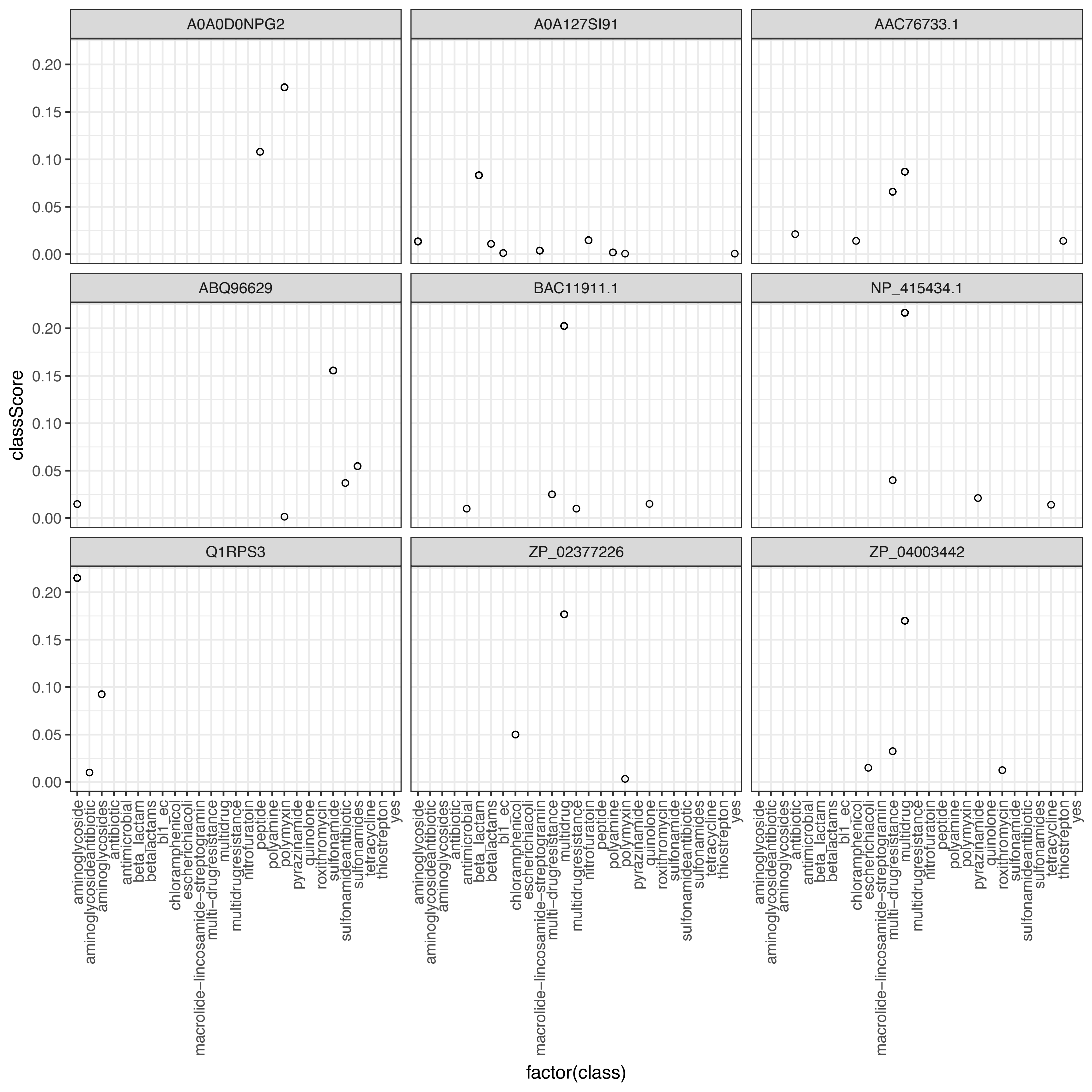

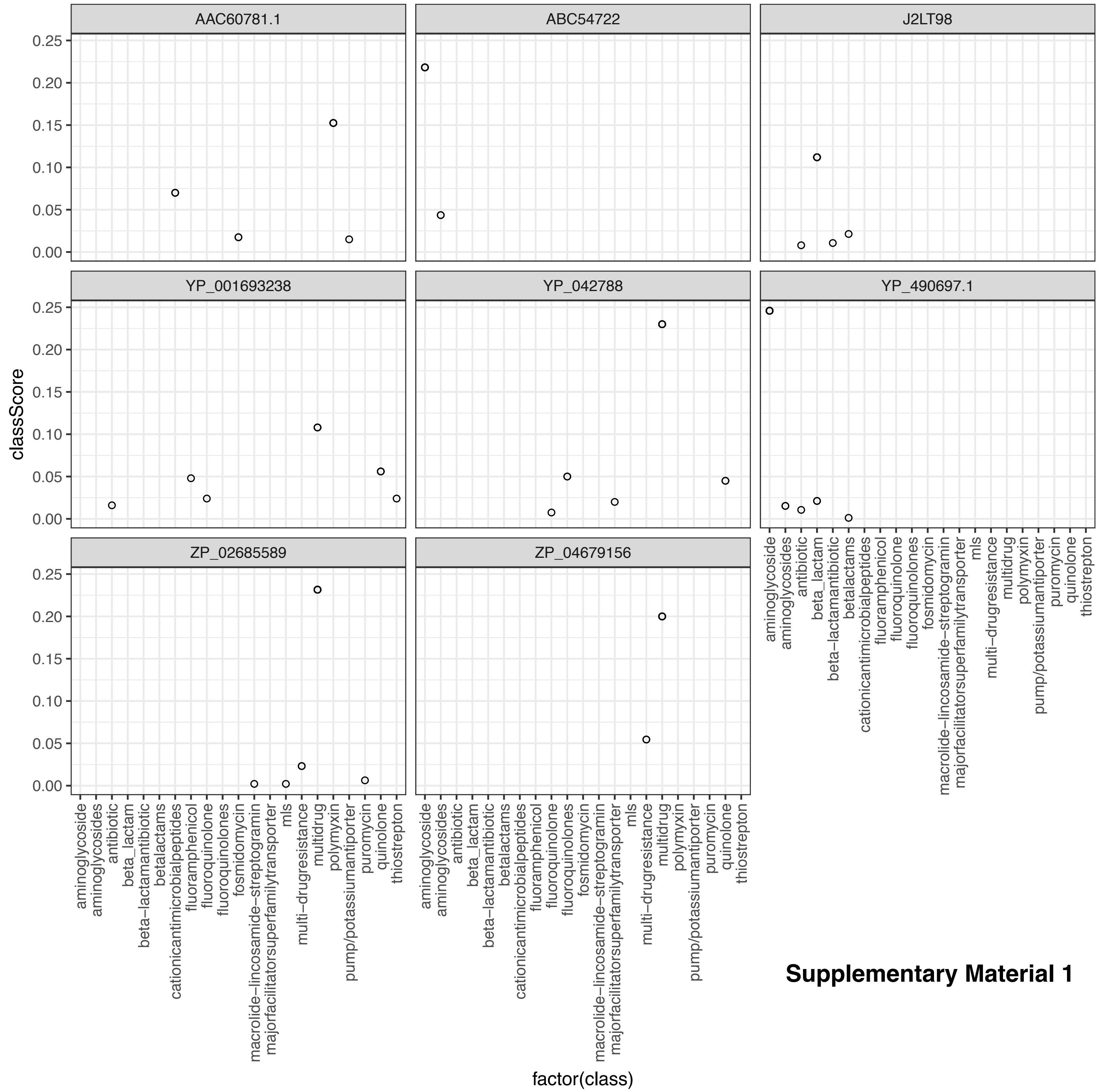


**Supplementary Material 1**

### SF 2

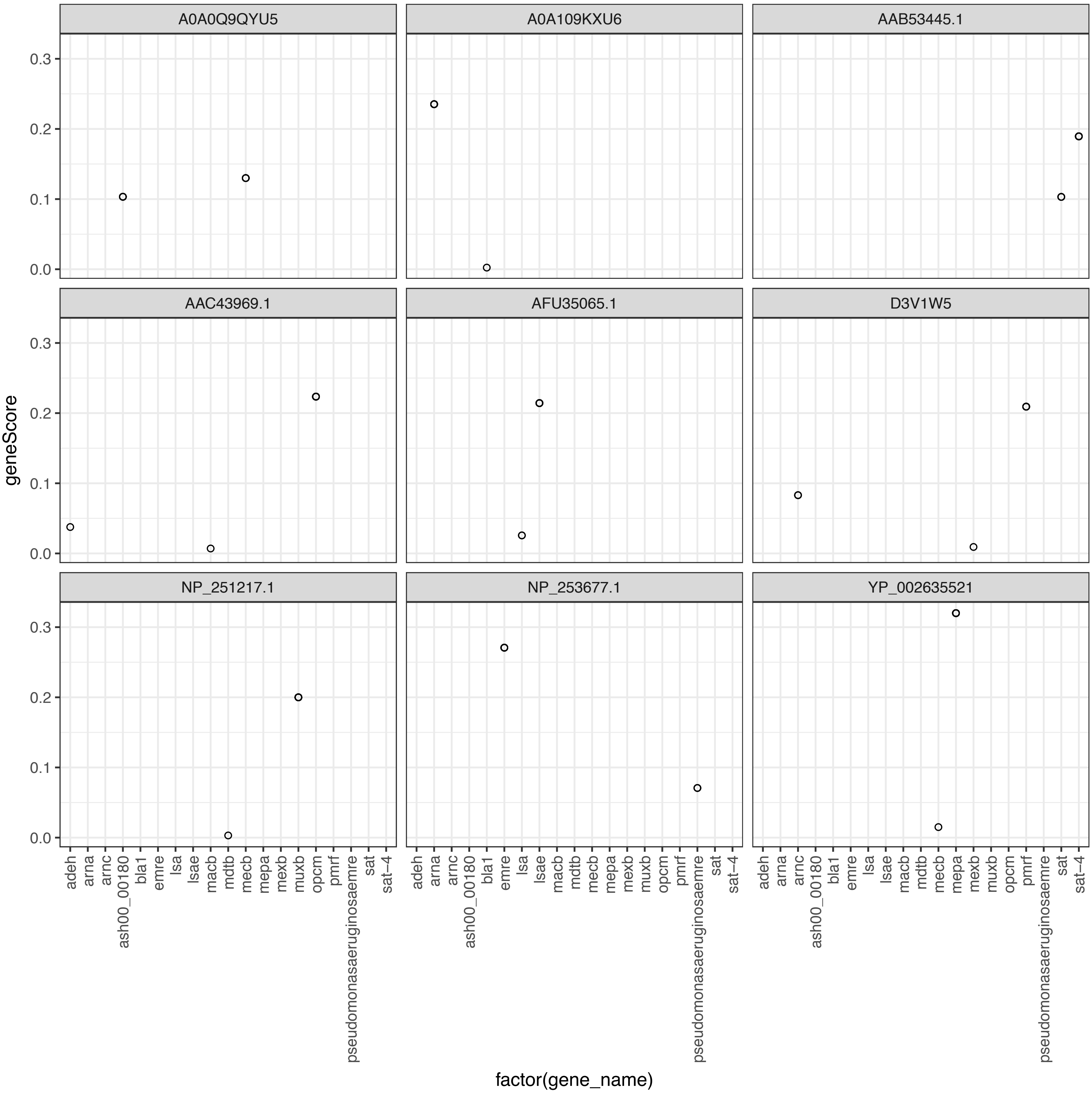

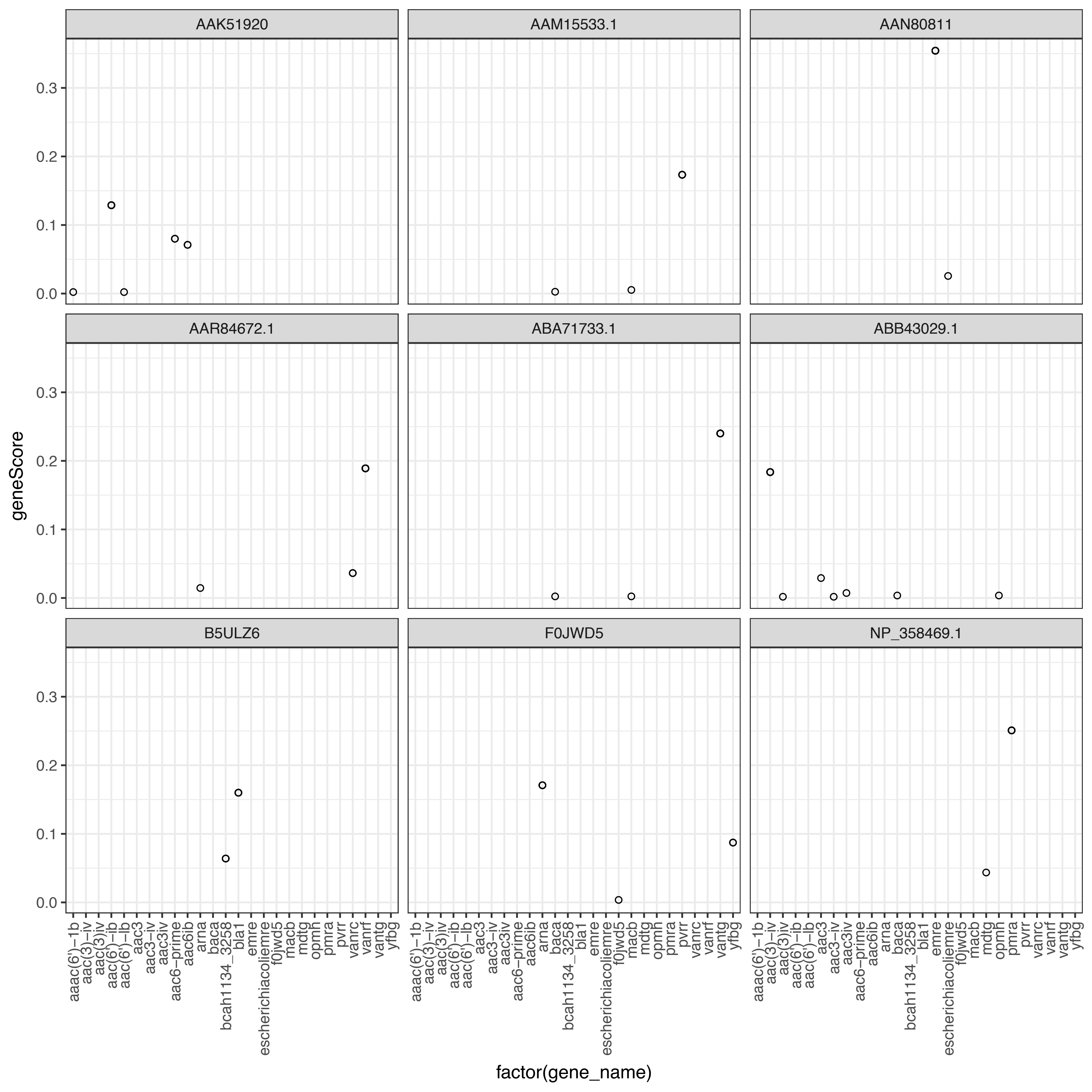

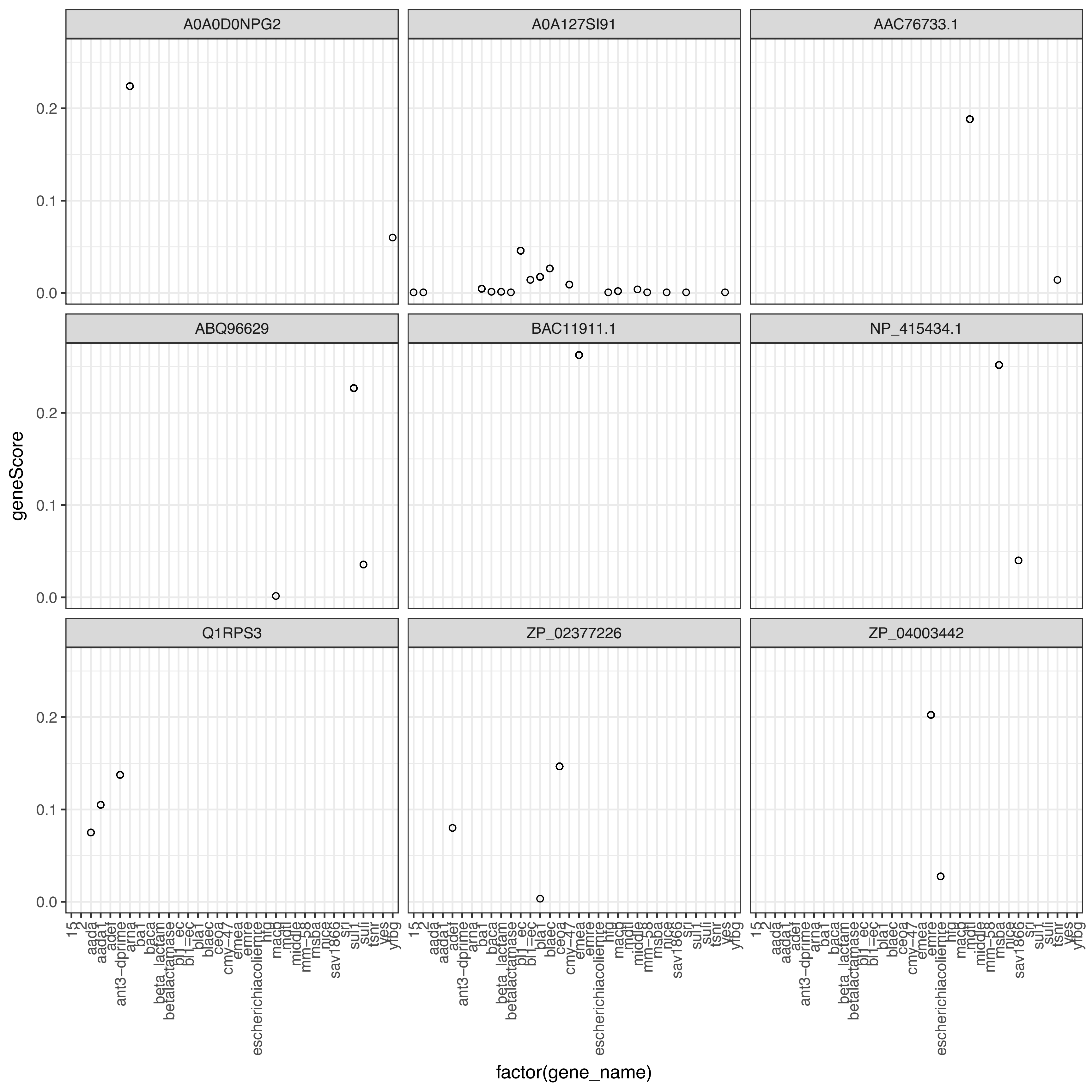

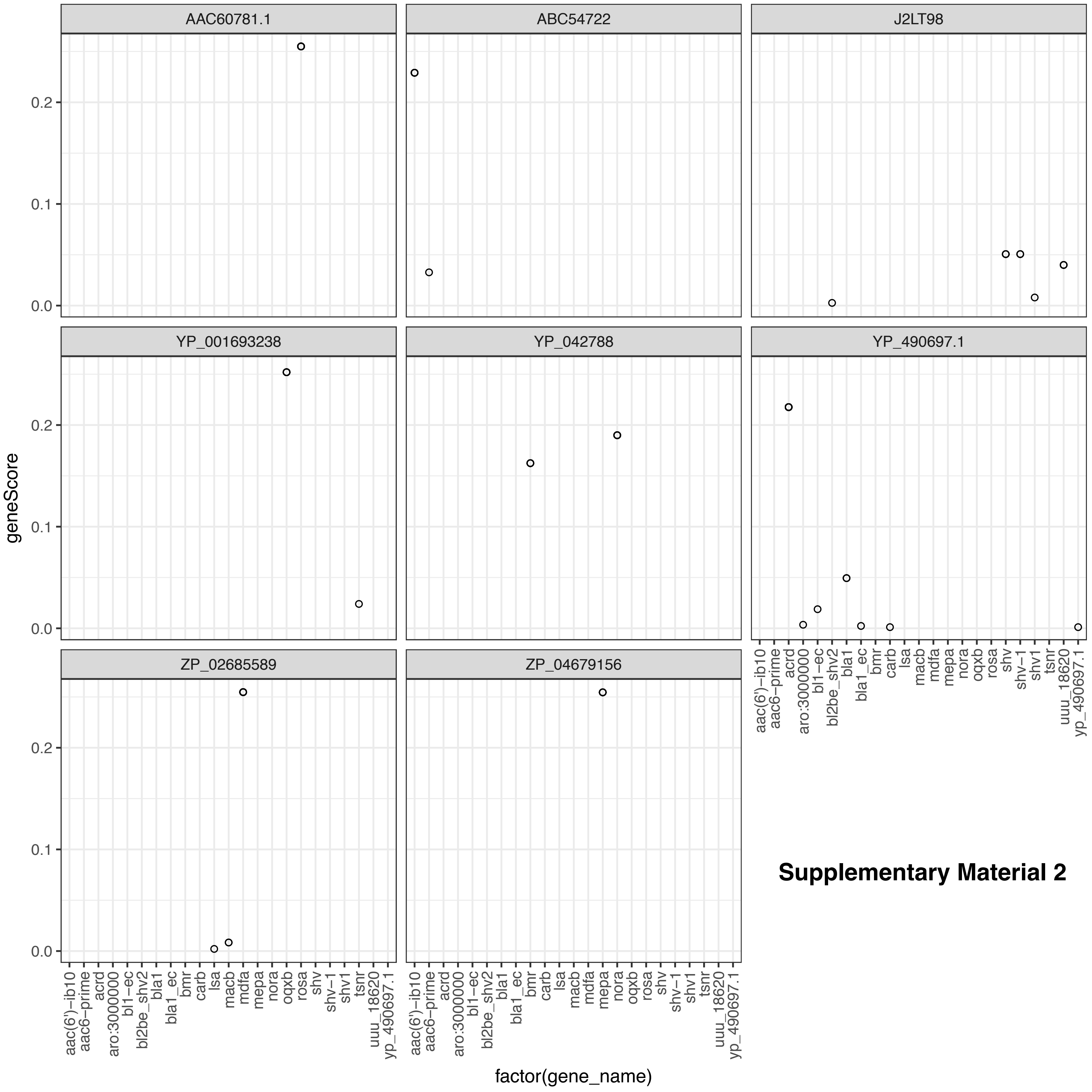


**Supplementary Material 2**
